# Trimester-Dependent Vertical Transmission of H5N1 Influenza Virus Through Placental and Mammary Routes Impairs Offspring Development

**DOI:** 10.1101/2025.07.07.663583

**Authors:** Brittany A. Seibert, Maclaine A. Parish, Sattya N. Talukdar, Tianle Zhang, Jennifer A. Liu, Sabal Chaulagain, Jaiprasath Sachithanandham, Oscar Hernandez, Cosette Schneider, Lynda Coughlan, Andrew S. Bowman, Richard J. Webby, Jack R. Harkema, Andrew Pekosz, Sabra L. Klein

## Abstract

Avian influenza H5N1 has pandemic potential and historically has caused more severe disease in pregnant women than the general population. With increasing transmission of H5N1 detected among placental mammals, animal models are necessary for testing countermeasures, including during pregnancy. Pregnant outbred mice were infected with a contemporary strain of bovine H5N1 during the second or third trimester equivalent. Second trimester infection caused *in utero* transmission, with infectious virus detected in the uterus, placenta, and fetus. Birth following third trimester infection resulted in offspring with decreased size, neurodevelopmental delays, and adolescent behavioral impairments, with infectious virus detected in the neonatal milk ring and lungs, as well as mammary tissues. H5N1 viral protein colocalized with trophoblast cells in the placenta and epithelial cells in mammary tissue that spatially overlapped with lectins for α2,3-linked SA. With the pandemic potential of H5N1, our vertical transmission model in placental mammals is essential for understanding viral spread and evaluating treatments during pregnancy.

## Introduction

Pregnancy represents a distinct physiological and immunological state, which causes more severe outcomes from influenza infections than in non-pregnant individuals^1,2^. Pregnant women have historically been at a greater risk of severe disease and death following highly pathogenic avian influenza (HPAI) infection, as well as seasonal and pandemic influenza A virus (IAV)^1,2^. A recent analysis of 30 pregnant women diagnosed with avian influenza virus infection (H5N1, H5N6, and H7N9) reported that more than half (n=16) were infected with H5N1, which occurred across gestational ages, with all but two resulting in maternal and *in utero* fetal death^2^. A prior study reported the presence of viral RNA (vRNA) in both placental and fetal tissues of pregnant mice infected with H5N1(A/Shenzhen/406H/2006) IAV, though viral-cell tropism and infectious virus within these tissues, as well as maternal morbidity, were not evaluated^3^. Despite poor maternal and infant outcomes of HPAI H5N1 infections, there is limited information on viral pathogenesis during pregnancy and its impact on the developing offspring.

As avian influenza viruses continue to evolve, there has been a dramatic increase in mammalian infections of H5N1 viruses from clade 2.3.4.4b, with notable spread among numerous placental mammals, including cattle^4–6^. While multiple transmission routes (i.e, oral, ocular, mammary, aerosol, fomites) have been examined using strains from the recent cattle outbreak^7–10^, vertical transmission of HPAI influenza infection during pregnancy remains poorly studied. Emerging reports now suggest possible vertical transmission of H5N1 clade 2.3.4.4b among South American pinnipeds, including cases with placental lesions and viral RNA (vRNA) and protein detected in both placental and fetal tissues^11^, alongside reports of aborted fetuses collected during an H5N1 outbreak in Uruguay^12^ and Argentina^13^. Cattle farms have also reported that H5N1-infected heifers exhibit reproductive complications, including abortions during mid- to late-gestation^14,15^. Continued circulation of H5N1 among dairy cattle, along with spillover into other mammalian species, increases the risk of human exposure and underscores the need to better understand the impact of HPAI H5N1 infection during pregnancy and the potential for vertical transmission.

We established a pregnancy mouse model to study the pathogenesis and transmission dynamics of a contemporary bovine H5N1 IAV strain from the 2024 outbreak. We intranasally inoculated pregnant CD-1 outbred mice at two gestational ages to investigate both *in utero* and post-birth transmission to offspring. Lastly, we evaluated the impact of maternal H5N1 infection on birth outcomes, neonatal growth, and postnatal neurodevelopment. Our study provides the groundwork for assessing H5N1 infection during pregnancy by establishing a pregnant mouse model that can be extended to study other HPAI virus strains and provide insights into viral dissemination, vertical transmission dynamics, and long-term offspring outcomes.

## Results

### Bovine H5N1 influenza A virus dissemination and pathogenesis in non-pregnant and pregnant female mice

We examined the pathogenesis of two B3.13 genotype bovine isolates, A/bovine/Texas/98638/2024 (H5N1; A/bovine/TX) and A/bovine/Ohio/B24OSU-439/2024 (H5N1; A/bovine/OH) in non-pregnant outbred female mice. We utilized outbred mice because they better approximate the tolerogenic immune responses required to prevent rejection of semi-allogeneic fetuses^16^, and exhibit improved survival following pandemic H1N1 challenge compared with inbred strains^16–18^. We intranasally inoculated non-pregnant females with 100 TCID50 of A/bovine/TX, A/bovine/OH, or media. Similar morbidity (weight change and rectal temperature) was observed for both viruses (**Fig. 1a-b**). Peak clinical signs of disease appeared earlier in mice infected with A/bovine/TX (5 days post-infection [dpi]) compared to those infected with A/bovine/OH (9 dpi) (**Fig. 1c**). All mice infected with A/bovine/TX succumbed by 6 dpi, while those infected with A/bovine/OH succumbed by 9 dpi (**Fig. 1d**). At 5 dpi, females infected with A/bovine/TX exhibited greater infectious viral loads in the nasal turbinates (NT), mammary glands, ovaries, and nervous system tissues (spinal cord, olfactory bulb, frontal lobe, and hindbrain) than females infected with A/bovine/OH (**Fig. 1e**). Similar viral loads were detected in whole blood, trachea, lungs, and colon. We selected A/bovine/OH for subsequent experiments in pregnant mice to increase the probability of survival. When we infected non-pregnant mice with a lower dose of A/bovine/OH, although not statistically significant, mice infected with 10 TCID50 showed reduced morbidity and mortality compared to those infected with 100 TCID50 (**Fig. 1f-g**).

**Fig. 1:**
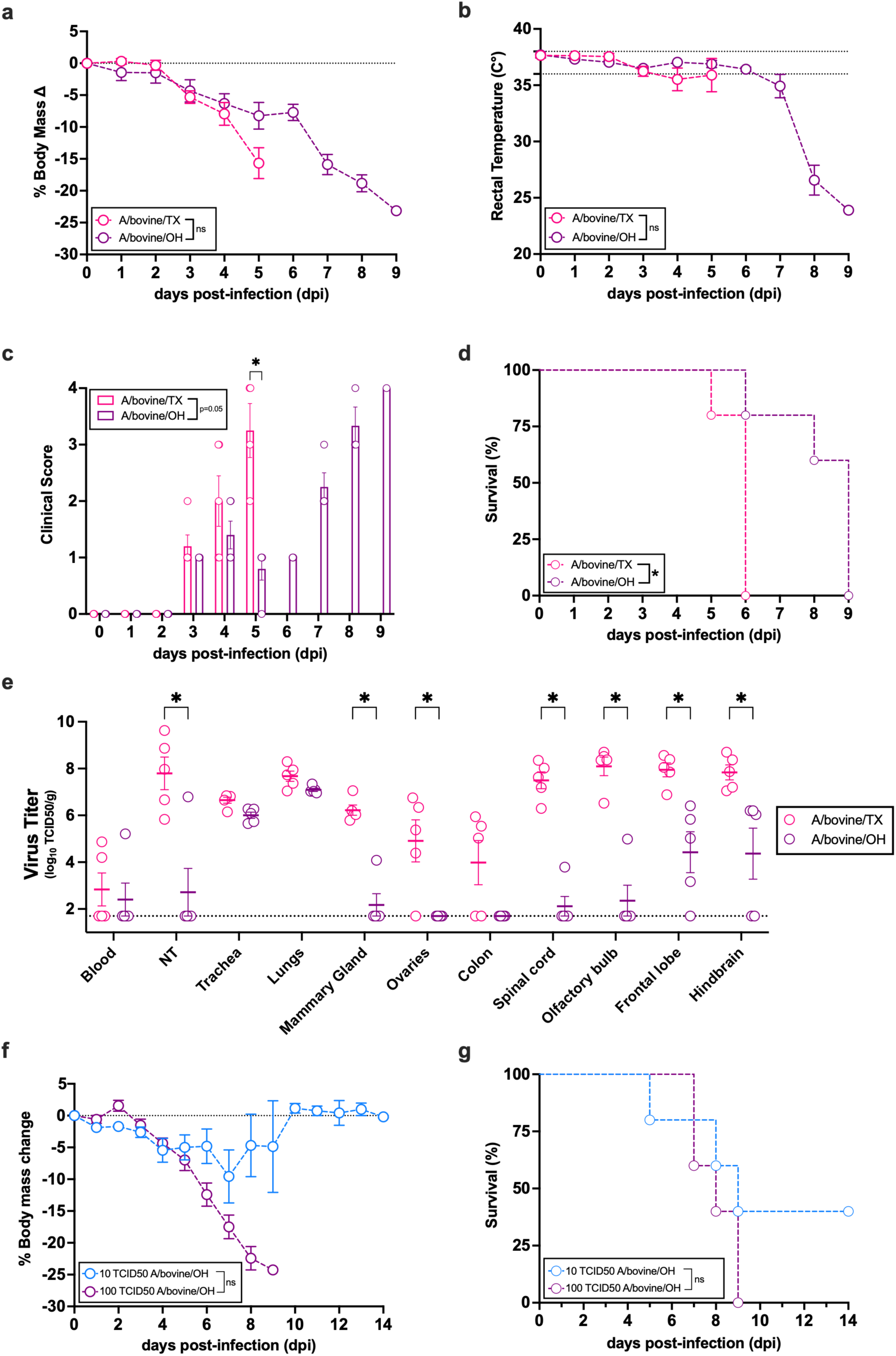
Pathogenicity of bovine H5N1 IAV in nonpregnant female CD-1 mice. CD-1 female mice (7-8 weeks, n=5/group) were intranasally inoculated with 100 TCID50 of A/bovine/TX or A/bovine/OH. (**a**) Body mass, (**b**) rectal temperature (dashed line shows normal temperature range), (**c**) clinical scores (scale of 0–4), and (**d**) survival were recorded daily. (**e**) At 5 dpi, mice inoculated with A/bovine/TX or A/bovine/OH (7-8 weeks, n=5/group) were euthanized and tissues were collected for viral quantification by TCID50 assay. **(f, g)** CD-1 female mice (7-8 weeks, n=5/group) were inoculated with 10 or 100 TCID50 of A/bovine/OH. (**f**) Body mass and (**g**) survival were monitored daily for 14 days. (a-b, f-g) Individual symbols or (c) bars represent the mean ± standard error of the mean (SEM) per group, with (e) individual mice indicated by symbols. Asterisks represent significant differences (p ≤ 0.05) performed using a (a-c,f-g) two-tailed unpaired t-test of the calculated area under the curve (AUC), (c) multiple unpaired t-test, (d) Mantel-Cox, or a (e) two-way ANOVA with Tukey post-hoc test.

To assess whether pregnancy increases morbidity and viral dissemination, we intranasally inoculated 10 or 100 TCID50 of A/bovine/OH into non-pregnant or pregnant (dams) mice at embryonic day (E) 10, roughly corresponding to the human second trimester^16,19^. Dams infected with 100 TCID50 exhibited less weight gain through 6 dpi than dams infected with 10 TCID50, a dose-dependent effect that was not observed in non-pregnant females (**Fig. 2a**). Rectal temperature indicated that inoculation dose, more than pregnancy status, influenced morbidity (**Fig. 2b**), in which both pregnant and non-pregnant mice given 100 TCID50 showed greater hypothermia compared to those given 10 TCID50.

**Fig. 2:**
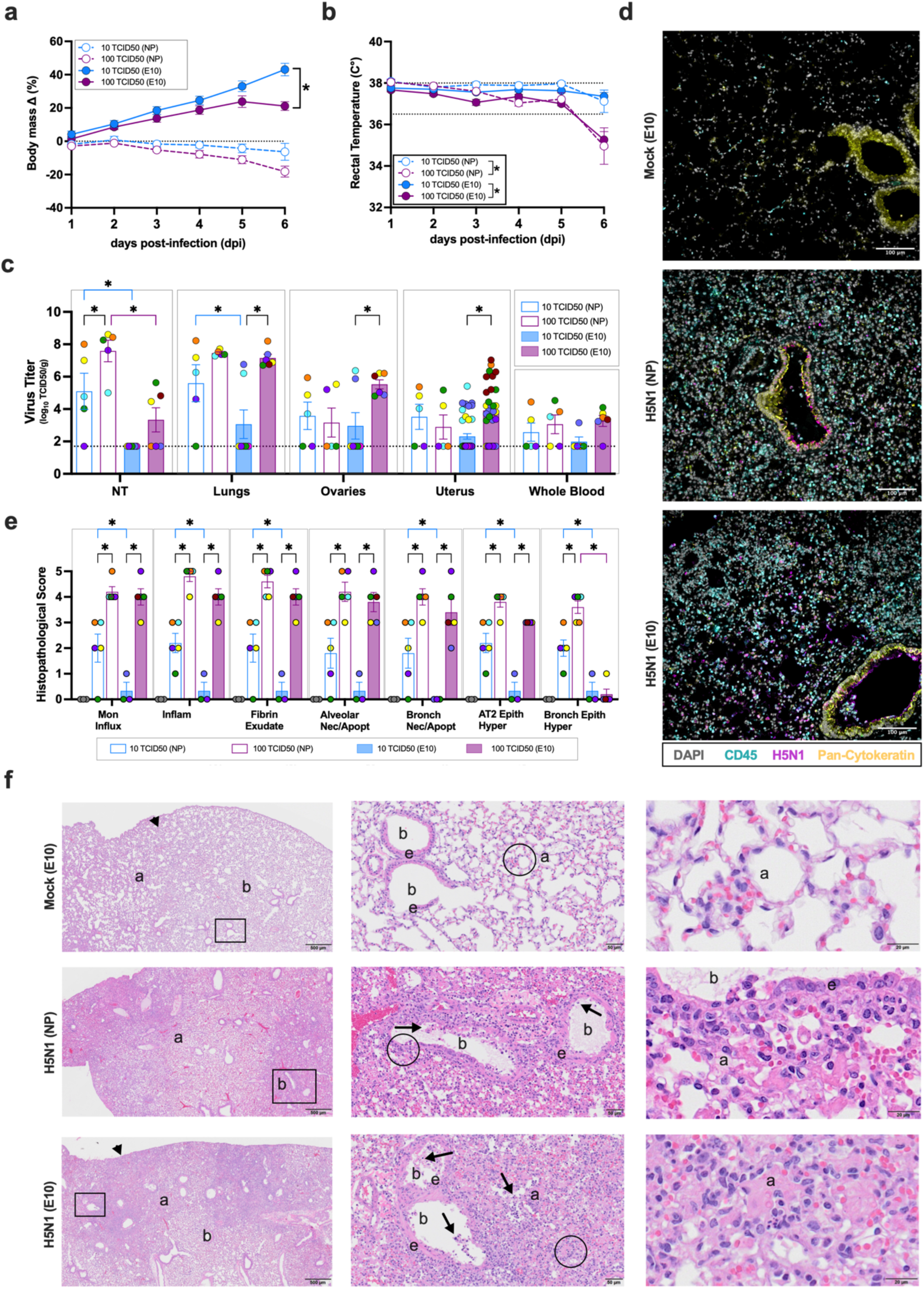
Pathogenesis of bovine H5N1 IAV in non-pregnant and pregnant female mice. CD-1 non-pregnant (NP; 7-8 weeks; n=5/group) or pregnant (E10; n=6-7/group) female mice were intranasally inoculated with 10 or 100 TCID50 of A/bovine/OH. Mock controls were inoculated with media. (**a**) Body mass change and (**b**) rectal temperature (dashed line shows normal temperature range) were recorded daily. (**c-e**) At 6 dpi, tissues were collected (n=5-7/group). (**c**) Infectious virus was quantified by TCID50 assay in the nasal turbinates (NT), lungs, ovaries, and whole blood. Individual points are color-coded by animal, while bar colors indicate group. (**d**) Immunofluorescent staining of fixed lung tissue. Cell nuclei (DAPI) are labeled in grey, CD45 in cyan, pan-cytokeratin in yellow, and H5N1 virus protein in magenta. Images were captured at 20X (scale bar: 100 µm). (**e**) Severity of individual lesions (histopathology) in lung tissue from NP and E10 dams was scored by a board-certified pathologist for monocyte influx (Mon Influx), inflammation (Inflam), fibrin exudate, alveolar necrosis and apoptosis (Alv Necr Apopt), bronchiolar necrosis/apoptosis (Bronch Necr/Apopt), AT2 epithelial hyperplasia (AT2 Epith Hyper), and bronchiolar epithelial hyperplasia (Bronch Epith Hyper). (**f**) Light photomicrographs of hematoxylin and eosin–stained lung tissue captured at different magnifications (low (scale bar: 500 µm), medium (scale bar: 50 µm) and high (scale bar: 20 µm)) from right to left, respectively. Squares and circles indicate the location of magnified images to the right. Arrows in the lower middle photo highlight virus-induced inflammatory cell exudate in airway and alveolar airspaces─a, alveoli and b, bronchiole. Arrows in the right lower photo highlight the presence of apoptotic epithelial cells and apoptotic bodies lining alveolar septa, with accumulation of mixed inflammatory cells and proteinaceous fluid in alveolar lumens. Individual symbols (a) or bars (b,d) represent the mean ± SEM per group, with individual mice indicated by the color-coded points (b,d). Asterisks represent significant differences (p ≤ 0.05) determined by a repeated measures two-way ANOVA (a) or two-way ANOVA with Tukey post-hoc test (b,d) and colored by comparisons (black, inoculum dose (10 vs. 100 TCID50 within NP or E10); blue, 10 TCID50; purple, 100 TCID50 (NP vs. E10).

The tissue distribution of infectious virus revealed that non-pregnant females had greater viral loads in the NT and lungs at 6 dpi (**Figure 2C**). Dams, particularly those receiving 100 TCID50, showed greater detectable infectious virus in the ovaries and uterus. Infectious virus was detected in the ovaries and in at least one uterine horn in 6/6 dams (100 TCID50), compared to the ovaries and/or uterine horns of only 2/5 non-pregnant mice.

Histological examination of the lungs in non-pregnant and pregnant females infected with 100 TCID50 of A/bovine/OH revealed viral protein localized in luminal epithelial cells lining distal preterminal and terminal bronchioles and centriacinar alveoli, coinciding with an influx of CD45 positive immune cells that had spatial overlap of viral protein (**Fig. 2d**). This is consistent with reported immune cell tropism of a North American 2.3.4.4b H5N1 IAV^20^. Co-staining of viral protein with lectins that recognize sialic acid (SA) linkages α2,3-gal-β(1-4) on N- and O-linked glycans (MAL-I), SA α2,3-gal-β(1-3) on O-linked glycans (MAL-II), and SA α2,6-gal/GalNAc (SNA)^21^, revealed a greater number of H5N1 positive cells also positive for MAL-I and MAL-II compared to SNA (**Supplementary Fig. 1**). As expected, pulmonary histopathology was greater in pregnant mice infected with 100 TCID50 compared to those infected with 10 TCID50 or mock-infected (**Figure 2E**). Non-pregnant females experienced greater pulmonary pathology than pregnant females after infection with 10 TCID50, suggesting greater pulmonary susceptibility to disease (**Fig. 2e**). While no histopathology was present in the lungs of mock-infected mice, infected females had moderate to severe pulmonary histopathology characterized by a locally extensive to widespread subacute necrotizing alveolitis and bronchiolitis (**Fig. 2f**). Infected non-pregnant and pregnant lungs (100 TCID50) had virus-induced apoptosis, necrosis, and exfoliation of airway epithelium lining distal, small-diameter bronchioles extending to terminal bronchioles in the centriacinar regions of the lung. Bronchiolar epithelial lesions were accompanied by an acute to subacute mixed inflammatory cell infiltrate composed of neutrophils and mononuclear cells (lymphocytes and monocytes) along with intramural and luminal accumulation of proteinaceous fluid (**Fig. 2f, arrows**). Epithelial and inflammatory cell lesions extended into alveolar ducts and alveoli characterized by a necrotizing and exudative alveolitis with scattered focal areas of apoptotic epithelial cells and apoptotic bodies lining alveolar septa, and accumulation of mixed inflammatory cells and proteinaceous fluid in alveolar airspaces (**Fig. 2f**). Taken together, pregnancy results in less severe pulmonary tissue damage, but enhanced viral tropism in reproductive tissues, as compared with non-pregnant females.

### *In utero* vertical transmission of bovine H5N1 influenza A virus

To determine whether *in utero* vertical transmission occurred, placental and fetal tissues were examined for infectious virus at 6 dpi. Among dams infected with 10 TCID50 of A/bovine/OH, 2/7 litters (11 of 42 conceptuses [comprising the uterus, placenta, and fetus]) had detectable infectious virus in the placenta, with one litter also showing corresponding viral detection in the fetuses (4 of 42 conceptuses) (**Fig. 3a**). Dams infected with 100 TCID50 had infectious virus detected in 6/6 litters within the placenta (31 of 33 conceptuses), with three litters also exhibiting infectious virus in the fetuses (13 of 34 conceptuses), suggesting dose-dependent increased virus prevalence in the placenta and transmission to fetal tissues.

**Fig. 3:**
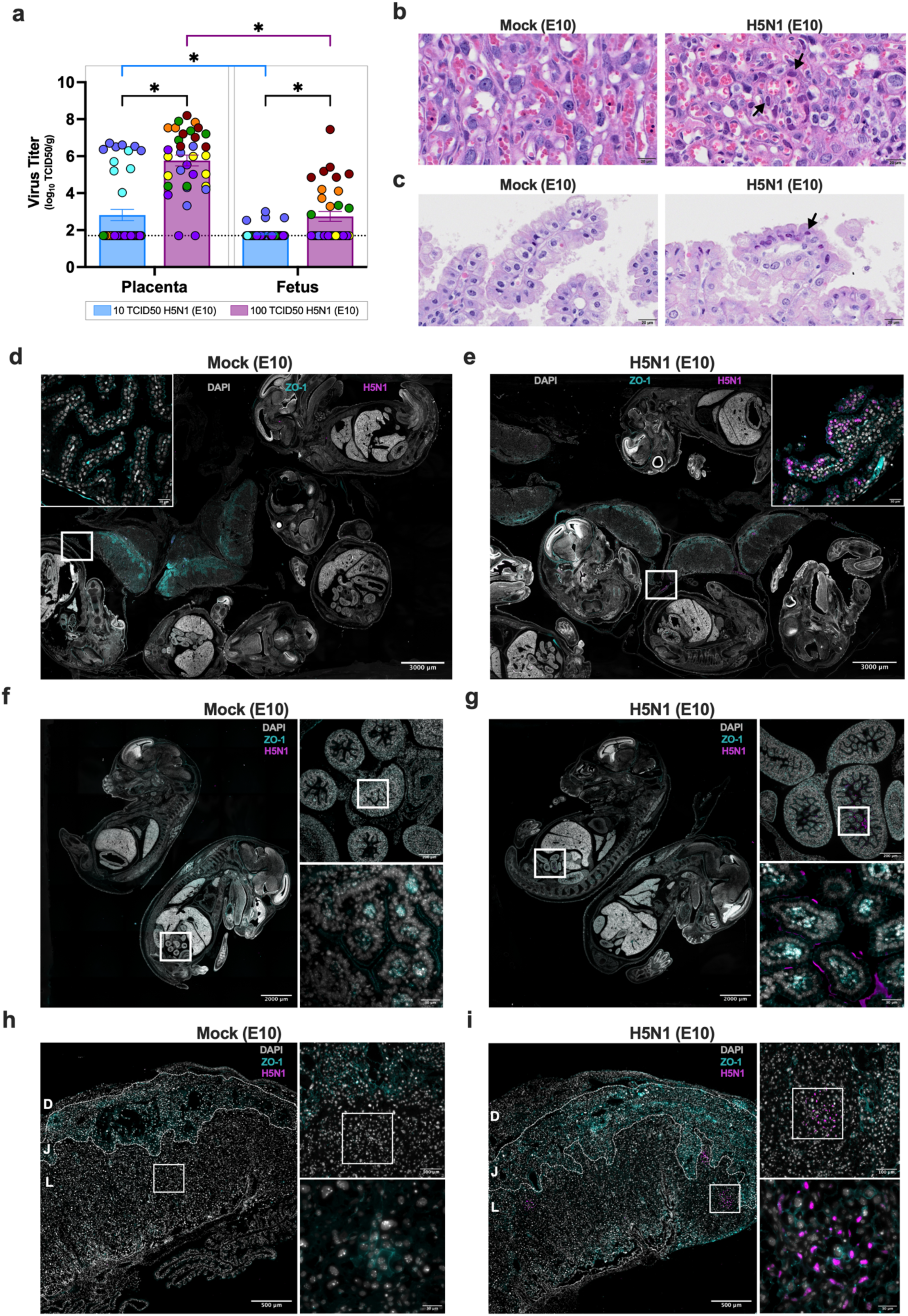
*In utero* vertical transmission and conceptus tissue tropism of bovine H5N1 IAV. CD-1 pregnant (E10) mice were intranasally inoculated with media, 10 or 100 TCID50 of A/bovine/OH. (**a**) At 6 dpi, dams (n=5-7/group) were euthanized, and infectious virus was quantified by TCID50 assay in a subset of placental and fetal tissues (n= 4-6/dam). The dashed line indicates the limit of detection. Individual points are color-coded by litter corresponding to the color of the dam in Fig. 2, while bar colors indicate viral dose. Hematoxylin and eosin (H&E)– stained tissue of the (**b**) placental labyrinth (mock or H5N1 (100 TCID50)) or (**c**) yolk sac (mock or H5N1 (10 TCID50)). Arrows represent virus-associated, small focal lesions. Immunofluorescent staining of the (**d-e**) conceptus, (**f–g**) fetus, and (**h-i**) placenta from E10 dams inoculated with (**d,f,h**) media or (**e,g,i**) A/bovine/OH (100 TCID50). The left images were taken at (**d-g**) 4X tiled images (scale bar: 2,000 or 3,000 µm) or (**h-i**) 4X (scale bar: 500 µm). Higher-resolution images were captured at (**d-e**; top right) 40X (scale bar: 30 µm) or (**f-i**; top right) 20X (scale bar: 100 or 200 µm) and (**f-i**; bottom right) 60X (scale bar: 30 µm). White boxes indicate the regions displayed in the magnified images. Cell nuclei (DAPI) are shown in grey, tight junctions (ZO-1) in cyan, and H5N1 virus protein in magenta. In panels h-i, the zones of the mouse placenta (the decidua (D), junctional zone (J), and labyrinth (L) are distinguished by dashed white lines and labeled accordingly. Bars (a) represent the mean ± SEM per group, with individual mice indicated by colored points. Asterisks represent significant differences (p < 0.05) by a two-way ANOVA with Tukey post-hoc test (a-b).

No histopathology was present in the placenta of mock-infected dams (**Fig. 3b**). For the embryonic age at the time of tissue collection (E16), there was an expected amount of degeneration and subsequent necrosis in the maternal decidua in mock dams and only slightly more decidual and junctional zone deterioration in H5N1-infected dams. In a subset of placentas from infected dams, virus-associated, small focal lesions were observed scattered within the labyrinth layer (**Fig. 3b, arrows**). These lesions consisted of trophoblasts with nuclear inclusions or apoptotic cell death features, e.g., pyknosis, karyorrhexis, apoptotic bodies and phagocytic cells containing cellular debris. Virus-associated histopathological changes were observed in the columnar epithelial cells lining the yolk sac (**Fig. 3c, arrow**).

Immunofluorescence staining for H5N1 antigen was utilized to identify the spatial distribution of the virus. We evaluated the distribution of viral protein in fixed conceptuses to examine viral dissemination in tissues disrupted during the dissection process. Viral antigen was absent in mock-infected dams (**Fig. 3d**), but was detected in the yolk sac of a conceptus from one infected dam (**Fig. 3e**). Because the yolk sac plays a key role in fetal nutrition^22^, we evaluated placental and fetal mass at 6 dpi. Maternal H5N1 infection at E10 results in significantly decreased placental but not fetal mass (**Supplementary Fig. 2a-b**). Because infectious virus was detected in a subset of fetuses without growth restriction at either dose, we examined the localization of viral protein in fixed fetuses from litters that exhibited vertical transmission. This can provide a more comprehensive evaluation, as fetal orientation in fixed conceptuses can obscure the sagittal plane. Viral protein, absent in fetuses from mock-infected dams (**Fig. 3f**), was localized in the intestinal tissue of fetuses from dams infected with either 10 (**Supplementary Fig. 2c**) or 100 (**Fig. 3g**) TCID50. With viral protein detected in the yolk sac and intestine of fetuses, we next examined the localization of viral protein in fixed placentas. The mouse placenta is composed of maternal decidua and fetal-derived compartments, including the junctional and labyrinth zones, where nutrient and gas exchange occur^23^. Viral protein was not observed in placentas collected from mock-infected dams (**Fig. 3h**); however, placentas from dams infected with 10 (**Supplementary Fig. 2d**) or 100 (**Fig. 3i**) TCID50 displayed viral protein localized to the decidua, junctional zone, and labyrinth zone, consistent with the distribution of virus-associated lesions noted in the labyrinth zone via histopathology.

We next evaluated the cellular tropism of bovine H5N1 IAV in the placenta. A limited number of CD45 positive cells colocalized with H5N1 viral protein within the junctional zone but not in the labyrinth, suggesting that immune cells were not the primary target in the placenta (**Fig. 4a**). Vimentin staining for endothelial and fibroblast populations^24^ revealed limited colocalization with H5N1 viral protein in the junctional and labyrinth zones (**Fig. 4b**). Trophoblasts are a major cell type within the junctional and labyrinth zones, comprising multiple populations with distinct functions essential for maintaining a healthy pregnancy^23,24^. H5N1 viral protein colocalized widely with pan-cytokeratin positive trophoblasts in the junctional and labyrinth zones (**Fig. 4c**). Co-staining of viral protein and pan-cytokeratin with lectins revealed that MAL-I was predominantly detected in the labyrinth zone, colocalizing with pan-cytokeratin positive cells and viral protein (**Fig. 4d**), while MAL-II exhibited very limited staining in the placenta (**Supplementary Fig. 3**). SNA was observed in both the junctional and labyrinth zones; however, most viral protein-positive cells did not colocalize with SNA (**Fig. 4e**). These findings indicate that maternal infection with bovine H5N1 IAV results in vertical transmission to the developing fetus, with trophoblasts as the predominant placental cell type that colocalizing with H5N1 viral protein and α2,3-linked SA.

**Fig. 4:**
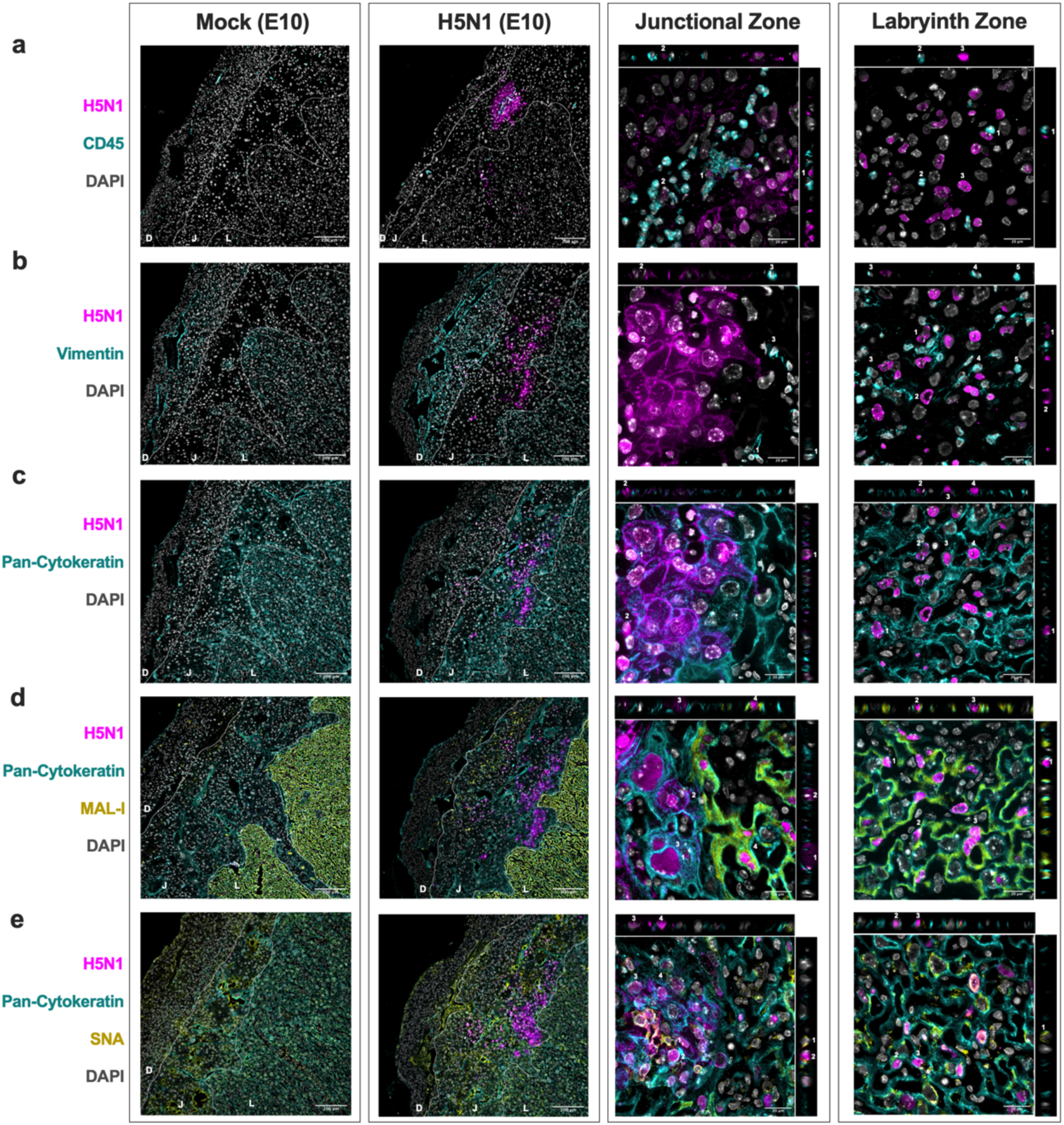
Cell tropism of bovine H5N1 IAV within the junctional zone and labyrinth zones of the mouse placenta. CD-1 pregnant (E10) mice were intranasally inoculated with media (Mock-E10), or 10 TCID50 of A/bovine/OH (H5N1-E10). At 6 dpi, dams (n=3-5/group) were euthanized and placentas (n=2-3/dam) were collected for histopathology analysis. Images in panels labeled Mock (E10) and H5N1 (E10) were captured at 10X (scale bar: 200 µm), while the right two panels labeled Junctional Zone and Labyrinth were taken at 100X (scale bar: 20 µm). In Mock and H5N1 (E10) panels, the three zones of the mouse placenta (the decidua (D), junctional zone (J), and labyrinth (L)) are distinguished by the dashed white lines and labeled accordingly. The Junctional zone and Labyrinth panels include orthogonal xz and yz views to assess colocalization of cell markers. The numbers identifying specific cells of interest correspond to those shown in the maximum projection at the center. Immunofluorescence staining of (**a**) CD45+, (**b**) vimentin+, or (**c**) pan-cytokeratin+ cells is displayed in cyan, cell nuclei (DAPI) are shown in grey, and H5N1 viral protein in magenta. (**d**) Immunofluorescence staining of pan-cytokeratin+ cells are displayed in cyan, MAL-I (α 2,3 SA) staining is shown in yellow, cell nuclei (DAPI) are shown in grey, and H5N1 viral protein is shown in magenta. (**e**) Immunofluorescence staining of pan-cytokeratin+ cells are displayed in cyan, SNA (α 2,6 SA) staining is shown in yellow, cell nuclei (DAPI) are shown in grey, and H5N1 viral protein is in magenta.

### Vertical transmission of bovine H5N1 influenza A virus post-birth

To investigate post-birth virus transmission, pregnant dams were inoculated with 10 or 100 TCID50 of A/bovine/OH during the human third trimester equivalent (E16)^16,19^. Dams, along with their entire litter, were euthanized at 1 or 4 days post-birth (dpb), corresponding to 4 or 7 dpi. At 4 dpi (1 dpb), only 2/5 litters from dams infected with 100 TCID50 had detectable virus in at least one of the pups’ lungs and /or milk ring, while virus was undetectable in all litters from dams infected with 10 TCID50 (**Fig. 5a**). We evaluated viral dissemination in milk and four distinct mammary glands^25^. At 7 dpi (4 dpb), a subset of dams infected with 10 (3/6) or 100 (3/6) TCID50 had detectable virus in their lungs and mammary tissue (**Fig. 5b**). Infectious virus was detected in at least one of the pups’ lungs and/or milk ring from 1/6 dams infected with 10 TCID50 and 2/6 dams infected with 100 TCID50. Pups with detectable infectious virus were born to dams that also had virus detected in the milk and thoracic, abdominal, and inguinal mammary tissue.

**Fig. 5:**
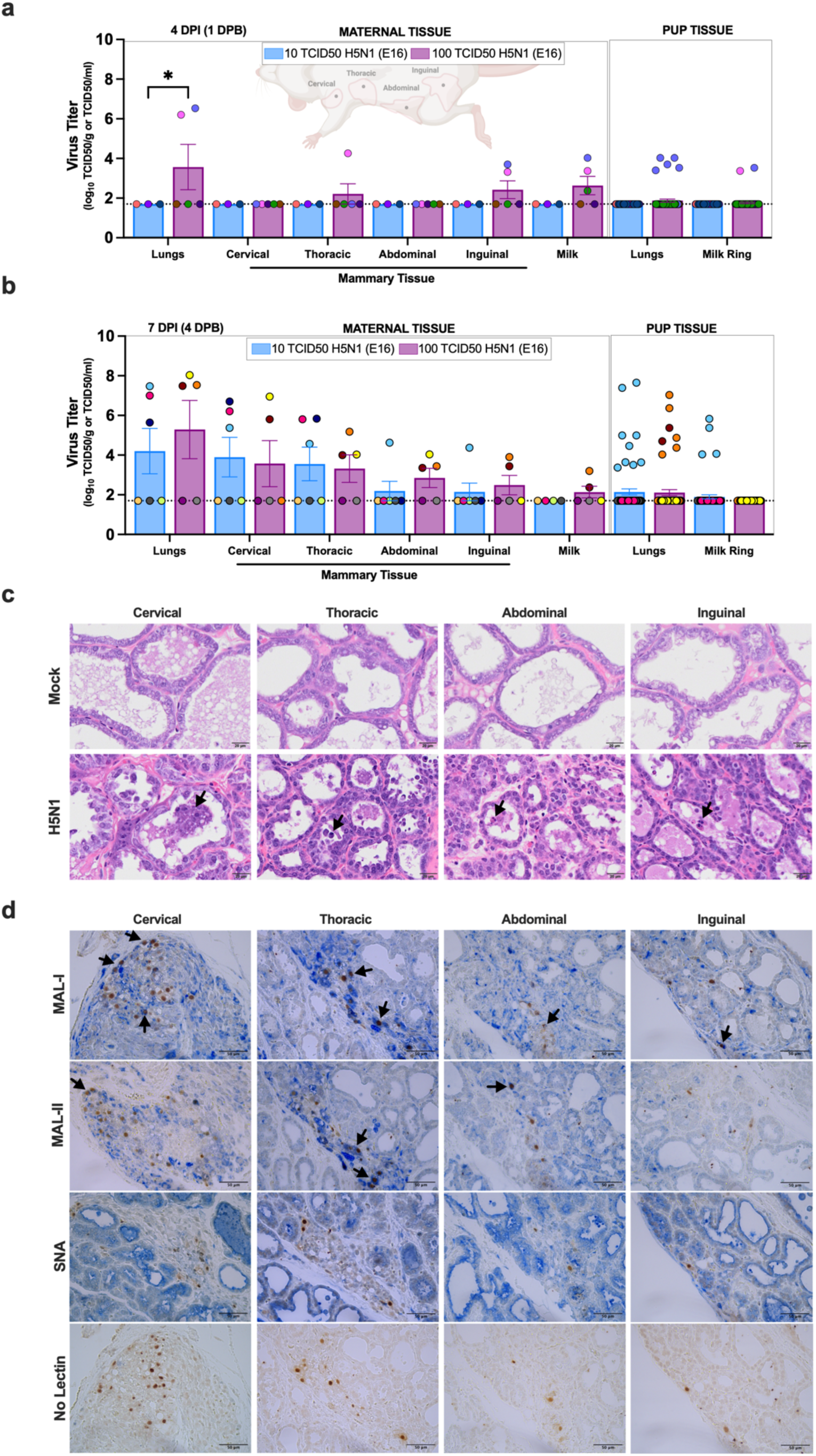
Post-birth vertical transmission and tissue tropism of bovine H5N1 IAV. CD-1 pregnant (E16) mice were intranasally inoculated with 10 or 100 TCID50 of A/bovine/OH. At (**a**) 4 dpi (1 dpb) or (**b**) 7 dpi (4 dpb), a subset of dams and their entire litters were euthanized. Infectious virus was quantified by TCID50 assay in the lungs, milk, and mammary tissue (cervical, thoracic, abdominal, and inguinal) of lactating dams, as well as in the pup lungs and milk ring. Individual points are color-coded by litter. (**c**) H&E–stained mammary tissue sections collected 7-8 dpi from E16 pregnant dams (mock – left and 100 TCID50 – right). Arrows indicate pyknotic or karyorrhectic nuclei. (**d**) Mammary tissue from E16 dams (100 TCID50) was stained for viral N1 antigen (brown chromogen), individually duplexed with MAL-I (α 2,3 SA), MAL-II (α 2,3 SA), or SNA (α 2,6 SA) staining. A control lacking the lectin primary antibody is included. Arrows indicate H5N1-positive cells. Images were captured at 40X (scale bar: 20 µm). Illustration depicting various mammary glands was created in BioRender. Seibert, B. (2025) https://BioRender.com/tq8un5s. Bars (a-b) represent the mean ± SEM per group. Asterisks represent significant differences (p < 0.05) by a two-way ANOVA with Tukey post-hoc test.

We determined whether the presence of H5N1 in mammary tissues resulted in pathological changes at 7 dpi (4 dpb). No histopathology was present in any of the lactating mammary glands of mock-infected dams (**Fig. 5c**). Terminal acinar lumens in the mock-infected dams contained a mixture of lipid and proteinaceous material, and the epithelial cells lining the lumens exhibited marked cytoplasmic vacuolation due to the presence of secretory lipid droplets. The terminal acinar lumens in the mammary glands of H5N1-infected dams were noticeably less distended, containing proteinaceous material, and exfoliated epithelial cells, with some cells exhibiting pyknotic or karyorrhectic nuclei (**Fig. 5c, arrows**). Basophilic regenerative epithelium predominantly lined the acinar lumens of the H5N1-infected mice. When examining the presence of viral antigen co-stained with lectins by immunohistochemistry, cytoplasmic and nuclear staining of the N1 protein was observed in the epithelial cells lining the terminal acinar lumens, with co-staining predominantly observed with MAL-I and MAL-II lectins (α2,3-linked SA) in mammary tissue from dams infected with 100 TCID50 (**Fig. 5d, arrows**), but not mock-infected dams (**Supplementary Fig. 4**).

Compared to dams infected with 10 TCID50, those infected with 100 TCID50 exhibited increased morbidity, along with increased clinical signs of disease during the first week post-infection (**Fig. 6a-b**). To further evaluate vertical transmission at later time points during lactation, dams were inoculated with 10 TCID50 A/bovine/OH at E16 to increase survival. Infected dams did not exhibit changes in morbidity, but clinical signs of disease peaked at 9 dpi (**Fig. 6a-b**). Between 7 and 9 dpi, virus was detected in the lungs, mammary glands, and milk in a subset of dams (**Fig. 6c**), whereas no infectious virus was detected in tissues collected from dams euthanized at 28 dpi (**Fig. 6d**). Tissues from subsets of pups within individual litters were examined for infectious virus 1-25 dpb, which corresponded to 4-28 dpi. While no pups had detectable virus at 1 dpb (4 dpi), virus was detected in the milk ring, lungs, and blood from pups at 4 dpb (7 dpi) and 6 dpb (9 dpi) (**Fig. 6e**). In contrast, no virus was detected in pups at either 12 dpb (15 dpi) or 25 dpb (28 dpi).

**Fig. 6:**
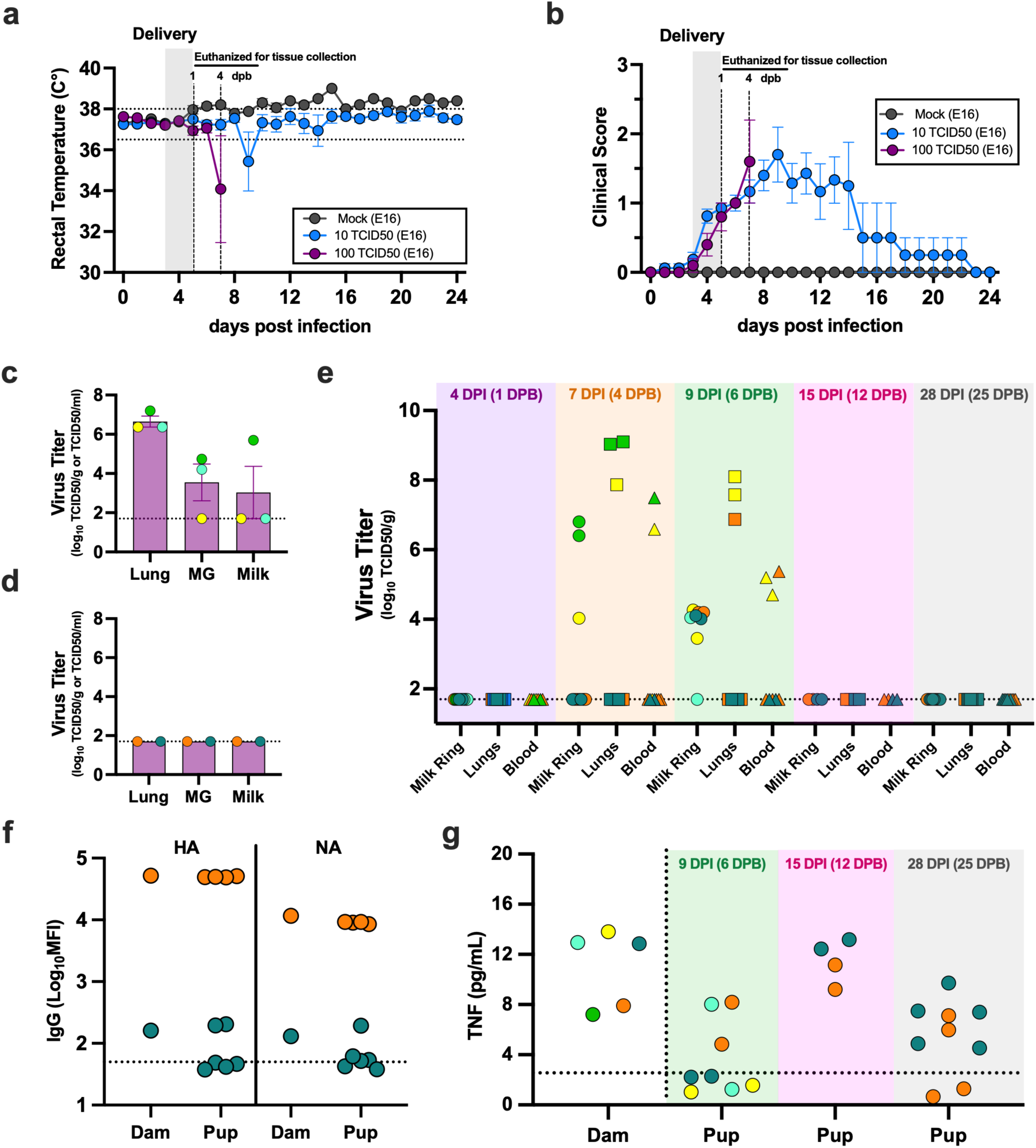
Post-birth bovine H5N1 IAV transmission dynamics to offspring. CD-1 pregnant (E16) mice were intranasally inoculated with media or 10 or 100 TCID50 of A/bovine/OH. (**a**) Rectal temperature (dashed line shows normal temperature range was recorded daily). (**b**) Clinical scores (scale of 0–4) were documented daily. The shaded grey bar marks birth. Dams inoculated with 100 TCID50 were euthanized at 4 or 7 dpi, indicated by a dashed line, while a separate group of mock-infected and 10 TCID50^-^infected dams was monitored up to 24 dpi. Between (**c**) 4-6 and (**d**) 28 dpi, a subset of dams (n=2-3/dpi) inoculated with media or 10 TCID50 of A/bovine/OH were euthanized, and infectious virus was quantified by TCID50 assay in the lungs, mammary glands (MG), and milk. (**e**) A subset of pups (n= 4-10/dpb) euthanized at 4 days dpi (1 dpb), 7 dpi (4 dpb), 9 dpi (6 dpb), 15 dpi (12 dpb), and 28 dpi (25 dpb) and the milk ring, lung, and whole blood, were collected and analyzed for infectious virus by TCID50 assay. H5N1 HA and NA specific (**f**) IgG were measured in serum from dams and their corresponding pups euthanized at 28 dpi (25 dpb) using systems serology. The results are reported as log_10_ mean fluorescence intensity (MFI). (**g**) Spleens collected from dams and their corresponding pups at 9 dpi (6 dpb), 15 dpi (12 dpb), or 28 dpi (25 dpb) were examined for TNF using a ProcartaPlex Cytokine kit. Individual points are color-coded by individual dam and their corresponding pups. Individual points (a-b) or bars (c-d) represent the mean ± SEM per group. The dashed line indicates the limit of assay detection (c-e) or background (f-g).

To confirm that dams and pups at 25 dpb (28 dpi) were infected, antibody responses to H5N1 were measured. H5- and N1-specific IgG titers were detectable in both dams euthanized at 28 dpi (**Fig. 6f**). While the pups from the dams euthanized at 28 dpi did not have infectious virus, a subset of pups had detectable H5- and N1-specific IgG titers, suggesting either transfer of maternal antibodies or possible infection (**Fig. 6f**). To evaluate whether maternal infection induced inflammatory immune responses in dams and their offspring, TNF concentrations were measured in spleens from dams at either 7-9 dpi or 28 dpi and offspring at 6, 12, or 28 dpb (**Fig. 6g**). Splenic TNF was highly detectable in all dams at 7-28 dpi and in a subset of offspring at 6, 12, or 25 dpb, suggesting that maternal H5N1 infection induced acute and persistent inflammation in dams and pups, even after virus clearance. Taken together, bovine H5N1 IAV caused acute infection in dams infected during the third trimester, resulting in postnatal vertical virus transmission likely through lactation, with virus clearance but persistent inflammation in both the dam and the pups 28 days post-infection.

### Postnatal outcomes and long-term neurocognitive impairments in offspring following maternal bovine H5N1 influenza A virus infection

Because maternal and offspring systemic inflammation (i.e., TNF) persisted following acute infection, we evaluated developmental and cognitive behaviors in the offspring. To determine adverse developmental outcomes from maternal H5N1 infection, litters from dams infected with 10 or 100 TCID50 of A/bovine/OH at E16 were evaluated. Infection with either dose did not impact litter size (**Supplementary Fig. 5a**) or cause pre-term birth (**Supplementary Fig. 5b**). Bovine H5N1 IAV infection of dams resulted in growth restriction in pups, as indicated by their smaller birth weight (**Fig. 7a**), shorter body length (**Fig. 7b**), and smaller head diameter (**Fig. 7c**). Because dams infected with 100 TCID50 were euthanized by 7 dpi due to increased morbidity (**Fig. 6a–b**), neurodevelopmental assessments were only conducted on pups from dams infected with 10 TCID50. At 5 and 9 dpb, cliff aversion (**Supplementary Fig. 5c and 7d**), measuring vestibular function, negative geotaxis (**Supplementary Fig. 5d and 7e**), measuring sensorimotor competence and vestibular function, and surface righting (**Supplementary Fig. 5e and 7f**), measuring motor development^16,26^, were delayed in offspring of H5N1-compared to mock-infected dams.

**Fig. 7:**
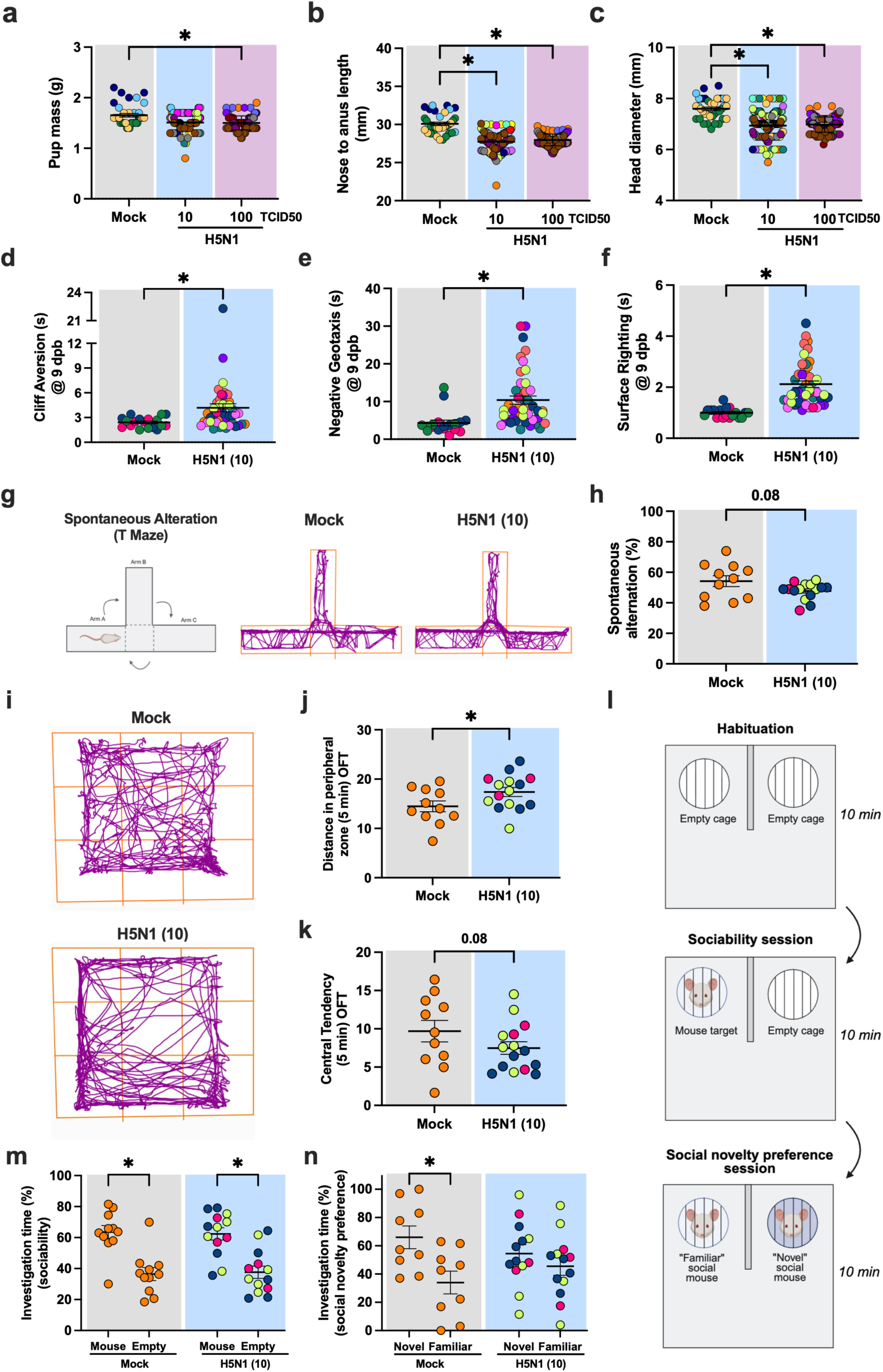
Perinatal outcomes following bovine H5N1 IAV infection during the third-trimester of gestation. CD-1 pregnant (E16) mice were intranasally inoculated with media or 10 or 100 TCID50 of A/bovine/OH. Within the first 12 hours of birth, (**a**) body mass, (**b**) length, (**c**) and head diameter of each pup were measured. (**d-f**) At 9 dpb, a subset of pups from dams inoculated with media or 10 TCID50 of H5N1 was assessed for (**d**) cliff aversion, (**e**) negative geotaxis, and (**f**) surface righting. (**g**) At 22 dpb, cognitive function was assessed using a spontaneous alternation task in a T-maze, as illustrated in the BioRender diagram, Liu, J. (2025) https://BioRender.com/j3x92wj., along with representative activity track plots from offspring born to mock or H5N1-infected dams. (**h**) Spontaneous alternation was based on the number of three consecutive arm entries, calculated as the percentage of correct alternations from total possible alternations. (**i**) Locomotor and anxiety-like behaviors were assessed at 23 dpb using the open field test (OFT); representative activity track plots from pups born to mock or H5N1-infected dams. (**j-k**) Anxiety-like behavioral measures were assessed during the first 5 min of OFT, including **(j)** distance traveled in the peripheral zone of the OFT and (**k**) central tendency that was quantified by measuring the distance traveled in the central zone compared to the total distance traveled. (**l**) The sociability and social novelty preference test consists of three consecutive testing phases, each with a 10-min testing interval, encompassing habituation, sociability, and social novelty, as illustrated in the BioRender diagram. Liu, J. (2025) https://BioRender.com/j3x92wj. Social preference during the (**m**) sociability and (**n**) social novelty preference sessions is reported as a percentage of total investigation time between the novel and inanimate object/familiar mouse, per individual pup. Individual points are color-coded by litter, while shaded colors indicate group: grey for media, blue for 10, and purple for 100 TCID50-inoculated pregnant dams. Horizontal bars indicate mean values across samples, with bars indicating the SEM for each group. Asterisks represent significant differences (p ≤ 0.05) by a (a-c, m-n) two-way ANOVA with Tukey post-hoc test or (d-k) two-tailed unpaired t-test.

To investigate whether maternal H5N1 infection resulted in long-term behavioral abnormalities, after weaning, juvenile mice were assessed using a series of cognitive and social tests (**Supplementary Fig. 5f**)^27–29^. Spatial working memory was measured by quantifying spontaneous alternations in an adapted T-maze (**Fig. 7g**). Offspring born to H5N1-infected dams showed a trend toward reduced percentages of correct spontaneous alternation, suggesting potential long-term memory impairments (**Fig. 7h**). This was not caused by reduced arm entries (**Supplementary Fig. 5g**) or differences in exploratory activity during the test (**Supplementary Fig. 5h**) as compared with offspring from mock infected dams. Locomotor activity during open-field test did not differ between groups (**Supplementary Fig. 5i**); offspring born to H5N1-infected dams, however, exhibited increased anxiety-like behaviors, including greater distance traveled in the peripheral zone (**Fig. 7i-j**), a trend towards reduced central tendency (**Fig. 7k**), and increased time rearing in the first 5 min of the OFT (**Supplementary Fig. 5j**). Social behavior was first evaluated by comparing preferences between social target compared to an inanimate object (sociability session) followed by measuring social novelty preference (SNP) between novel or familiar mouse^30^ (**Fig. 7l**). Because investigation time differed between mice, particularly during the SNP (**Supplementary Fig. 5k-l**), social/object preference in both sessions was reported as the percentage of total investigation time. Offspring from both mock and H5N1-infected dams showed a preference for a novel mouse over an object (empty cage) in the sociability session (**Fig. 7m**). In the SNP session, pups from mock-infected dams exhibited a preference for novel mice as expected, whereas offspring from H5N1-infected dams interact significantly less with the novel mouse, showing no preference between social targets (**Fig. 7n**). These results were not influenced by an inherent side bias during the habituation session (**Supplementary Fig. 5m**), suggesting long-lasting impairments in aspects of social memory and behavior. H5N1 infection during pregnancy in mice results in both acute and persistent adverse fetal, perinatal, and juvenile outcomes, consistent with the intrauterine growth restriction observed in a human case of H5N1 infection^2,31^.

## Discussion

We developed a comprehensive model to study H5N1 IAV pathogenesis during pregnancy and observed distinct tissue and cell tropism, with elevated viral titers in the reproductive tissues of pregnant mice. We showed *in utero* vertical H5N1 IAV transmission following intranasal inoculation of outbred pregnant mice at E10, as indicated by infectious virus in placental and fetal tissues with viral protein localization in junctional and labyrinth zones, the yolk sac, and fetal intestines. H5N1 protein colocalized with trophoblast cells in the placenta and epithelial cells in mammary tissue that spatially overlapped with lectins for α2,3-linked SA. Dams infected at E16 also transmitted virus to pups post-birth, most likely via milk, resulting in both acute and long-lasting systemic inflammation and adverse perinatal and developmental outcomes. These results provide insights into the vertical transmission dynamics of H5N1 IAV and contribute to the risk assessment of a high-risk population.

Infection during the second trimester (E10) of pregnancy reflects case studies that analyzed tissues from H5N1-positive patients^2,31,32^. We observed reduced placental mass with minimal placental pathology and no significant change in fetal mass. Analysis of a woman at 4 months of gestation who succumbed to H5N1 infection revealed normal placental development corresponding to gestational age and no histopathological abnormalities in fetal tissue^32^. Acute necrotizing deciduitis, also observed in our mouse model, was identified in the placenta of the pregnant woman^32^. Although placental and fetal damage was minimal, the woman exhibited widespread pulmonary damage^32^, consistent with our findings. When examining vRNA or protein, the placental tissue from the pregnant woman showed infected cells in the chorionic villi, with vRNA in the fetal lungs and liver^32^. In our model, viral protein was detected in the junctional and labyrinth zones of the placenta and within the intestines of fetuses from H5N1-infected dams. Both the mouse labyrinth and human chorionic villi are sites of maternal-fetal gas and nutrient exchange and are considered homologous structures^33^. We hypothesize that vertical transmission may occur via the labyrinth zone or through fetal ingestion of virus-containing amniotic fluid, which peaks during mid-to-late gestation and is critical for fetal intestinal development^34^. A previous mouse study, which infected inbred pregnant mice with A/Shenzhen/406H/2006 (clade 2.3.4) at E10, reported detectable viral RNA in both placental and fetal tissues, but did not measure infectious virus or the spatial or cellular distribution of the virus in the placenta or fetus^3^. We show that infectious virus is detected in both the placenta and fetus, with minimal viral protein colocalization with immune or endothelial cells, but predominantly detected in trophoblasts in the placental junctional and labyrinth zones. The majority of virus-positive trophoblasts across both placental zones colocalized with α2,3-linked SA-binding lectins; a subset of trophoblasts, however, especially in the junctional zone, did not stain with the lectins tested, indicating the potential involvement of additional receptors that need to be further investigated. Postmortem analysis of the placenta from a pregnant woman with confirmed H5N1 infection revealed positive viral protein staining in Hofbauer cells and cytotrophoblasts, but not in syncytiotrophoblasts^32^. Although mice lack Hofbauer cells, in vitro studies have shown that synctiotrophoblasts can support virus replication of H1N1, H5N1, and H7N9^35^.Investigating viral dissemination and mechanisms of H5N1 *in utero* vertical transmission using animal models has implications beyond human health, extending to other mammalian species, including livestock and marine mammals.

Given reports of fatal systemic influenza in cats and mice following ingestion of unpasteurized colostrum or milk from H5N1-infected dairy cattle^9,10,36^, we also used this late-gestation infection model to investigate the transmission window of H5N1 IAV during lactation. Consistent with a previous study of H5N1 IAV-infected lactating BALB/c female mice^10^, we observed infectious virus in pups from infected pregnant mice at 7–9 dpi, corresponding to 4–6 dpb. Also consistent with the findings by Eisfeld et al., we observed detectable infectious virus in pups from 2 of 5 litters born to dams infected with 100 TCID50 at 7 dpi (4 dpb)^10^. In addition to its influence on human health, our data contribute to understanding the agricultural impact, as H5N1 was detected in neonatal goats in Minnesota^37^, lactating cattle in Texas^38^, and lactating sheep in the United Kingdom^39^. Virus replication has also been documented in the mouths of calves and can be transmitted to the mammary glands of lactating cattle, as well as suckling kids from H5N1-infected lactating goats^40^. In evaluating virus replication and associated pathological changes in mammary tissues, experimental intra-mammary infection in dairy cattle results in high viral loads in milk and infected udder quarters with limited local spread^41,42^. Viral antigen in the mammary tissue from inoculated quarters is detected in alveolar epithelial cells and is associated with alveolar shrinkage lined by swollen epithelial cells, and replacement of secretory alveoli with fibrous connective tissue^41^. This is consistent with our observation of viral antigen in the epithelial cells lining the secretory alveoli of infected mammary tissue following intranasal inoculation. Whether these pathological changes are due to virus-induced pathology or a consequence of decreased milk production leading to mammary tissue reduction requires further study. Together, our results and those of previous studies further emphasize the importance of understanding virus transmission during lactation to prevent bidirectional H5N1 spread between lactating mothers and their offspring.

To date, no studies have examined offspring development following maternal H5N1 infection, whether acquired *in utero* or postnatally. We observed that H5N1 infection during pregnancy results in intrauterine growth restriction and offspring with neurodevelopmental delays, in the absence of pre-term birth or changes in litter size. Offspring born to H5N1-infected dams exhibit long-term impairments, including impaired short-term spatial working memory and social behavior, and increased anxiety-like behaviors. While vertical transmission has not been observed following maternal H1N1 infection in mice, offspring from H1N1-infected dams showed similar neurodevelopment impairments^16^ and increased anxiety-like behaviors^43^. Epidemiological studies also show increased risk for schizophrenia in offspring of women exposed to influenza during pregnancy^44,45^. While it is unclear whether the offspring assessed for long-term neurocognitive deficits had an active virus infection at earlier timepoints, at 25 dpb, most pups had detectable splenic TNF, suggesting maternal and early-life immune activation, which can contribute to behavioral deficits^46^.

Mouse models of pregnancy are instrumental for uncovering mechanisms of disease pathogenesis and evaluating potential treatment options, particularly given the frequent exclusion of pregnant women from clinical trials^47^. While we established a mouse model of bovine H5N1 IAV infection during pregnancy, we observed differences in morbidity and tissue tropism between two H5N1 clade B3.13 strains (A/bovine/TX and A/bovine/OH). These findings highlight the need for further studies to elucidate the pathogenesis of diverse H5N1 genotypes (e.g., B3.13 vs. D1.1). Analysis of virus replication and dissemination at multiple timepoints during acute infection during the second trimester (E10) would also provide greater insight into the pathogenesis of H5N1. It is also unclear what selective pressure transmission via milk has on the virus itself, as this may differ significantly from selective pressure in the lungs or placentas. Evaluating changes in virus sequences in infected milk and in pups infected with virus-containing milk could help inform whether vertical transmission can lead to more efficient mammalian adaptation of H5N1 viruses. While the developed mouse model can provide the foundation for understanding H5N1 IAV pathogenesis during pregnancy, translation of these findings to humans should be approached with caution due to the distinct placental anatomical morphological differences in human and mouse pregnancies^1^. We have, however, used similar pregnancy models of SARS-CoV-2 and Zika virus infection to test the effectiveness of antivirals or immunomodulatory drugs at mitigating maternal and offspring morbidity, which is best accomplished in preclinical models^48,49^. While a recent study demonstrated that baloxavir improves disease outcomes compared to oseltamivir after a lethal challenge of bovine H5N1 in non-pregnant female mice, baloxavir is not currently recommended for use in pregnant women^50,51^. Therefore, this model can provide a valuable framework for evaluating the efficacy of antivirals administered during pregnancy in reducing not only virus titers but also maternal morbidity and downstream effects on fetal and offspring outcomes. Observations from this model provide the groundwork for understanding the vertical transmission of bovine H5N1 viruses in humans and other placental mammals, including cattle, where adverse pregnancy outcomes have recently been reported following H5N1 infection^14,15^.

## Methods

### Animal experiments

Adult (8–12 weeks of age), non-pregnant female or timed-pregnant CD-1 mice were purchased from Charles River Laboratories. Pregnant mice were received on embryonic day (E) 8 or E14. Five non-pregnant female mice were housed per cage, while each pregnant mouse was singly housed. Mice were housed under standard animal biosafety level 3 (ABSL3) housing conditions with *ad libitum* food and water in the ABSL3 facility at the Johns Hopkins School of Medicine. Mice were given at least 24 hours to acclimate to the ABSL3 facility before infection^16^. All monitoring and experimental procedures were consistently performed at the same time each day. Studies were conducted under Biosafety Level 3 (BSL-3) containment and approved by Johns Hopkins University Animal Care and Use Committee (ACUC; Protocol MO24H150).

### Cells

Madin-Darby canine kidney (MDCK) cells, a kind gift from Dr. Robert A. Lamb, were maintained in complete medium (CM) consisting of Dulbecco’s Modified Eagle Medium (DMEM, Sigma-Aldrich, St Louis, MO, USA) supplemented with 10% fetal bovine serum (FBS; Gibco, Waltham, MA, USA), 100 units/ml penicillin/streptomycin (Life Technologies, Frederick, MD, USA) and 2 mM Glutamax (Gibco, Waltham, MA, USA) at 37 °C and 5% CO2.

### Viruses

The viruses utilized in this study: A/bovine/Texas/98638/2024 (H5N1; A/bovine/TX) and A/bovine/Ohio/B24OSU-439/2024 (H5N1; A/bovine/OH) were kindly provided by Dr. Andrew Bowman (The Ohio State University, Columbus, OH, USA) and Dr. Richard Webby (St Jude Children’s Research Hospital, Memphis, TN, USA)^52^. Virus stocks of A/bovine/TX and A/bovine/OH were generated by infecting Madin-Darby canine kidney (MDCK) cells at a multiplicity of infection (MOI) of 0.01 infectious units per cell, at 33°C using infection media (CM with no FBS and 5 μg/ml of N-acetyl Trypsin) as previously described^53^.

### Inoculation and monitoring of mice

Mice were anesthetized via intraperitoneal injection of a ketamine/xylazine cocktail (80 mg/kg ketamine, 5 mg/kg xylazine) prior to inoculation. Mice were intranasally inoculated with either 10 or 100 TCID50 of A/bovine/Texas or A/bovine/Ohio in 30μL of DMEM. Following intranasal inoculation, body mass, rectal temperature, and appearance of clinical signs were recorded once daily in the morning until tissue collection (4-28 days post-infection [dpi]). Clinical scores were reported on a scale of 0–4, with 1 point assigned for each of the following: piloerection, dyspnea, hunched posture, and absence of an escape response, on each day^54,55^.

### Milk collection from lactating mice

Lactating dams were separated from their litters for at least 30 min to allow milk accumulation. Mice were then anesthetized via intraperitoneal injection of a ketamine/xylazine cocktail (80 mg/kg ketamine, 5 mg/kg xylazine) followed by intraperitoneal injection of oxytocin acetate salt hydrate (0.05 mg/kg; Sigma-Aldrich, St Louis, MO, USA) to stimulate milk production. Milk was then expressed from the teat through manual stimulation and collected using a pipette, as previously described^56^. Milk was pooled from all teats during the collection process. One pregnant dam infected with 10 TCID50 succumbed to the infection just before sample collection; therefore, milk was not available for virus titration.

### Tissue and serum collection from mice

Experimental pregnant and non-pregnant female mice were euthanized via ketamine/xylazine overdose (160 mg/kg ketamine, 10 mg/kg xylazine) followed by cardiac exsanguination. For dams infected at E10, tissues were collected at 6 dpi. For dams infected at E16, tissues were collected at 4, 7, 9, 15, and 28 dpi, unless dams succumbed to infection prior to the assigned collection date, then tissues were flash-frozen on dry ice one day prior. Half of the whole blood was saved for virus titration, while the remaining volume was centrifuged at 10,000 × g for 30 minutes at 4°C to separate the serum. For the collection of reproductive tissues in pregnant mice, the uterine horns were removed, and every other conceptus starting at the top of the left uterine horn was flash frozen (the placenta, fetus, and uterine tissue were separated and frozen individually). This was performed until a total of 6 of each tissue were collected per dam. For tissues collected from pups after birth, pups were anesthetized via isoflurane and euthanized via decapitation. Tissues were collected and flash-frozen on dry ice at the specified time points post-birth.

### Virus Titration

Frozen tissues were thawed and placed into Lysing Matrix D tubes (MP Biomedicals, Santa Ana, CA, USA) with homogenization media (500mL DMEM, 5mL penicillin/streptomycin) with a minimum volume of 400μL and a maximum volume of 1200μL (10% weight/volume). Tissues were homogenized at 4.0 m/s for 45 seconds in a MP Fast-prep 24 5G instrument (MP Biomedicals, Santa Ana, CA, USA and centrifuged at 500 *g* for 5 minutes. Supernatants were collected, aliquoted, and stored at –80°C until further analysis. The supernatant from tissue homogenates was titrated by TCID50 assay using the Reed and Muench method^53,57^.

### Histopathology

Selected tissues, including lungs, placentas, fetuses, conceptuses, and mammary tissues were collected and fixed in formalin (Z-FIX, Fisher Scientific, Waltham, MA, USA) for histopathological examination. Prior to fixation, the left lung was perfused with formalin. Conceptuses remaining after flash-freezing tissues for viral titration were fixed, with the placenta, fetus, and uterine tissue dissected and fixed individually, beginning at the top of the left uterine horn. When there were more than 12 conceptuses, whole conceptuses (fetal, placental, and uterine tissue) were fixed without separation of individual tissues. Tissues were fixed for at least 72 hours prior to paraffin embedding and processing for routine histopathology with hematoxylin and eosin staining (H&E). A board-certified pathologist, blinded to the study, subjectively scored tissues (n=3 mocks, n=4 10 TCID50, n=6 100 TCID50) as follows: (0) none; (1) mild; (2) mild to moderate; (3) moderate; (4) moderate to severe; (5) severe.

### Immunohistochemistry

An antibody targeting influenza A virus N1 (Goat; Polyclonal Anti-Influenza Virus N1 Neuraminidase (NA), A/New Jersey/8/1976 (H1N1), BEI Resources, Manassas, VA, USA; dilution 1:250) was used to perform immunohistochemistry on mammary sections. Briefly, tissues embedded in paraffin were dewaxed according to an established protocol^58^, and antigen retrieval was performed by incubating slides in citrate buffer (Abcam; Cambridge, UK) within a steamer for 45 min. Tissues were blocked with 3% hydrogen peroxide for 10 min, followed by an overnight block with bovine serum albumin (BSA; 5%) diluted in PBS-Tween (0.1%). The next day, tissues were blocked with ReadyProbes Streptavidin/Biotin Blocking Solution (ThermoFisher, Waltham, MA, USA) following the manufacturer’s instructions. Tissue sections were incubated with either MAL-I, MAL-II, or SNA lectins from Vector Laboratories (Newark, CA, USA; 1:100 dilution) for 1 hour at room temperature, washed, and incubated with Streptavidin, Alkaline Phosphatase conjugate (ThermoFisher, Waltham, MA, USA; 1:500 dilution) for 1 hour at room temperature. Lectin staining was visualized using the Vector Blue Substrate kit, Alkaline Phosphatase (Vector Laboratories, Newark, CA, USA), with Levamisole solution (Vector Laboratories, Newark, CA, USA) added to reduce endogenous alkaline phosphatase activity, as per the manufacturer’s instructions. Tissue sections were washed, permeabilized with PBS-Triton (0.3%) for 10 min, and then blocked with normal donkey serum (NDS; Jackson ImmunoResearch, West Grove, PA, USA; 10%) in PBS-Tween 20 (0.1%) for 10 min at room temperature. Tissue sections were incubated overnight at 4°C with the influenza A virus N1 antibody diluted in PBS-Tween 20 (0.1%) with NDS (3%), followed by washing with 0.1% PBS-Tween and incubation with Donkey anti-Goat IgG (H+L) Secondary Antibody, HRP (ThermoFisher, Waltham, MA, USA; 1:500 dilution) for 1 hour at room temperature. Antibody binding was visualized using DAB (3,3′-diaminobenzidine; Vector Laboratories, Newark, California, USA) as the chromogen following the manufacturer’s instructions. Tissue sections were mounted using FluorSave (Millipore Sigma; Burlington, MA) and imaged using a Nikon Eclipse Ti2-E microscope (Nikon, Minato City, Tokyo, Japan) at 40X and then enhanced using ImageJ^59^.

### Immunofluorescence

As in the immunohistochemistry protocol, paraffin-embedded tissues were dewaxed and subjected to antigen retrieval. Tissues were then permeabilized with PBS-Triton (0.3%), stained using TrueBlack Lipofuscin Autofluorescence Quencher in DMF (biotium, Fremont, CA, USA), and then blocked with normal goat serum (NGS; Jackson ImmunoResearch, West Grove, PA, USA; 5%) for 1 hour at room temperature. Slides were incubated overnight at 4°C with primary antibodies targeting ZO-1 (Mouse; ZO-1 Monoclonal Antibody (ZO1-1A12), ThermoFisher, Waltham, MA, USA; dilution 1:200), CD45 (Rat; CD45 Monoclonal Antibody (30-F11), Functional Grade, eBioscience, ThermoFisher, Waltham, MA, USA; dilution 1:100), Pan-Cytokeratin (Mouse; Cytokeratin Pan Antibody Cocktail, ThermoFisher, Waltham, MA, USA; dilution 1:200), Vimentin (Chicken; Vimentin Polyclonal Antibody; ThermoFisher, Waltham, MA, USA; 1:4000), and/or H5N1 (Rabbit; Rabbit Anti-H5 Serum Control Panel; BEI Resources, Manassas, VA, USA; dilution 1:200) diluted in 3% NGS. Tissue sections were washed with 0.1% PBS-Tween and incubated with Goat anti-Mouse IgG (H+L) Cross-Adsorbed Secondary Antibody, Alexa Fluor™ 488 (ThermoFisher, Waltham, MA, USA; dilution 1:1000), Goat anti-Mouse IgG (H+L) Cross-Adsorbed Secondary Antibody, Alexa Fluor™ 647 (ThermoFisher, Waltham, MA, USA; dilution 1:1000), Goat anti-Rabbit IgG (H+L) Cross-Adsorbed Secondary Antibody, Alexa Fluor™ 555 (ThermoFisher, Waltham, MA, USA; dilution 1:1000), Goat anti-Rat IgG (H+L) Cross-Adsorbed Secondary Antibody, Alexa Fluor™ 647 (ThermoFisher, Waltham, MA, USA; dilution 1:1000), and/or Goat anti-Chicken IgY (H+L) Secondary Antibody, Alexa Fluor™ 488 (ThermoFisher, Waltham, MA, USA; dilution 1:1000) for 2 hours at room temperature. Tissues were subsequently incubated with DAPI (ThermoFisher, Waltham, MA, USA; 1 μg/mL) for 4 min, as per the manufacturer’s instructions. Tissue sections were imaged as z-stacks using a Nikon Eclipse Ti2-E microscope (Nikon, Minato City, Tokyo, Japan) at 4X, 20X, and 60X (Figures 3D-I and S2A-C) or LEICA Thunder widefield microscope at 10X, 20X, and 100X (Figures 2D, 4, S1, S3). DAPI was further processed in LAS X using THUNDER technology, and orthogonal xz and yz views were generated within the LAS X software. Maximum-intensity projection images were subsequently merged and annotated using ImageJ^59^.

For immunofluorescent lectin staining, paraffin-embedded tissues were dewaxed and subjected to antigen retrieval as described. Tissues were stained using TrueBlack Lipofuscin Autofluorescence Quencher in DMF (biotium, Fremont, CA, USA), and blocked with BSA (5%) and NGS (5%) overnight at 4°C. The next day, tissues were blocked with ReadyProbes Streptavidin/Biotin Blocking Solution (ThermoFisher, Waltham, MA, USA) following the manufacturer’s instructions. Tissue sections were incubated with MAL-I or MAL-II from Vector Laboratories (Newark, CA, USA; 1:100 dilution) for 1 hour at room temperature, washed, and then incubated with Streptavidin, Alexa Fluor™ 594 Conjugate (ThermoFisher, Waltham, MA, USA; dilution 1:1000) or Sambucus Nigra Lectin (SNA, EBL) Fluorescein (Vector Laboratories, Newark, CA, USA; dilution 1:100). Tissue sections were washed, permeabilized with PBS-Triton (0.3%) for 10 min, and incubated with the primary antibody (Pan-Cytokeratin or H5N1) for 1 hour at room temperature. Tissue sections were washed with 0.1% PBS-Tween and incubated Goat anti-Rabbit IgG (H+L) Cross-Adsorbed Secondary Antibody, Alexa Fluor™ 488 (ThermoFisher, Waltham, MA, USA; dilution 1:1000) and Goat anti-Mouse IgG (H+L) Cross-Adsorbed Secondary Antibody, Alexa Fluor™ 647 (ThermoFisher, Waltham, MA, USA; dilution 1:1000) for 1 hour at room temperature. Tissues were subsequently incubated with DAPI (ThermoFisher, Waltham, MA, USA; 1 μg/mL) for 4 min, as per the manufacturer’s instructions. Tissue sections were imaged as z-stacks using LEICA Thunder widefield microscope at 10X and 100X. DAPI was further processed in LAS X using THUNDER technology, and orthogonal xz and yz views were generated within the LAS X software. Maximum-intensity projection images were subsequently merged and annotated using ImageJ^59^.

### Luminex Assays

Antibodies were measured through multiplex Luminex serological assays as previously described^60^. Antigens used for these analyses include Influenza A H5N1 (A/Texas/37/2024), A/dairy cow/Texas/24-008749-002-v/2024 Hemagglutinin HA Protein (SinoBiological, Beijing, China), and the bovine N1 construct was kindly provided by Lynda Coughlan (University of Maryland, MD, USA) and produced in Expi293F cells as previously described^61^. Each antigen was coupled to unique MagPlex bead regions (Luminex). Briefly, beads were activated by combining activation buffer (0.1M NaH2PO4; pH 6.2), 50mg/mL 1-ethyl-3-[3-dimethylaminopropyl] carbodiimide (EDC), and 50mg/mL Sulfo-NHS (N-hydroxysulfosuccinimide). Activated beads were coupled to 25 μg of antigen in coupling buffer (0.05 M MES, pH 5.0). Activated coupled beads were blocked with blocking buffer (PBS 1X, 0.1% BSA, 0.02% Tween-20, 0.05% sodium azide; pH 7.4), washed with washing buffer (PBS 1X, 0.05% Tween-20) and stored at 4°C in PBS. To measure immunoglobulin isotypes, serum was heat-inactivated at 56°C for 30min and diluted 1:25 in PBS. 5μL of diluted serum run in duplicate was incubated with 45μL of bead solution (20μL of each bead region in 20μL Luminex Assay Buffer per plate) in a 384-well plate format for 2 hours shaking at 750 rpm. Following incubation, the plates were washed three times with Procartaplex Wash Buffer. Isotypes were identified by 1 hour incubation shaking at 750 rpm with 45μL of PE-conjugated IgG antibody (Southern Biotech, Homewood AL, USA; IgG). Following final incubation, the plates were washed three times with Procartaplex Wash Buffer and resuspended in 50μL of Procartaplex Wash Buffer for reading on the Luminex Intelliflex. Samples were run on FlexMap 3D Low with a DD range of 7000-1700. For each assay, a minimum of 100 events was recorded per antigen and the MFI of secondary antibody-conjugated to PE for each bead region was measured to quantify antigen-specific immunoglobulin isotype responses in each sample. Limit of background (LOB) was set by the PBS negative control. Concentrations of TNF in spleen homogenates were quantified using a mouse Immune Procartaplex kit (ThermoFisher, Waltham, MA, USA) following the manufacturer’s instructions and read on a Luminex Intelliflex instrument.

### Offspring measurements and developmental assessments

Gestational age of delivery, litter size, and offspring from mock and H5N1-infected dams were measured within the first 12 h of birth. Measurements including pup mass (g), head size measured as ear-to-ear diameter (mm), and length defined as nose to anus (mm) were recorded for each pup. At 5 and 9 days after birth, a subset of pups was subjected to neurodevelopmental tests, including cliff aversion, negative geotaxis, and surface righting, as previously described ^16,26^. For each test, at least 2 males and 2 females were used to limit confounding litter effects. Each pup performed each test three times, and the best time was recorded for each. Tests were considered failed and assigned a time of 30 seconds if the task was not completed. For cliff aversion testing, pups were briefly placed with their forepaws on the edge of a box, and the latency to retreat from the ledge was recorded. Negative geotaxis was assessed by placing pups facing downward on a 45° incline and recording the time taken to rotate 180° to face upward. Surface righting was assessed by recording the time required for a pup to turn from a prone to an upright position.

### Behavioral testing

Behavior was assessed in experimental pups at 22 and 24 dpb in the ABSL-3 facilities, with all behavioral procedures conducted within the biosafety cabinet^27^. Behavioral tests were conducted, starting with the least aversive and progressing to the most aversive, and involved a combination of manual and automated scoring using video tracking software for analysis (ANYMaze, Stoelting). Experimental mice were habituated to the testing environment within the biosafety cabinet under similar testing conditions (light exposure, sound, vibrations) for a minimum of 30 min prior to testing. Biosafety cabinets and maze apparatuses were thoroughly cleaned with Vimoba, followed by 70% ethanol. Between individual experimental mice, the floor, walls, and testing objects were cleaned with 70% ethanol and wiped dry between trials to eliminate residual odors. Behavioral testing was conducted during the light phase due to biosafety constraints of testing mice in the ABSL-3 setting.

#### Spontaneous alternation T-Maze

Spontaneous alternation was assessed in a T-maze apparatus to measure short-term spatial working memory and cognitive function in experimental mice^27–29^. Experimental mice were placed in one out of three arms of the T-maze and left to freely explore for 6 min. Total number of arm entries, arm visit entry sequences, and locomotor activity patterns were tracked and recorded from video footage using ANYmaze behavioral software. Arm entries were defined as all four limbs crossing into the arm and percentage of correct alternation was calculated using the following formula: [(number of correct alternations / total possible alternations (total arm entries -2)) x 100].

#### Open Field Test (OFT)

Exploratory locomotor activity was assessed using an open field (OF) maze^27^. Experimental mice were placed in the testing apparatus and allowed to freely explore for 10 min. Total distance travelled and locomotor activity patterns were tracked and recorded from video footage using ANYmaze behavioral software. The OFT session was separated into two 5 min sessions to allow for assessments of anxiety-like behaviors in a novel environment (OF maze). The distance traveled within a central or peripheral zone of the OF apparatus was measured from video footage using Any Maze behavioral software. The number of rears/time spent rearing (seconds) was manually scored during the first 5 min of the test. Rearing was defined as experimental mice standing on their hind legs with both forepaws elevated in the middle of the maze or against a wall.

#### Sociability and social novelty preference test

Sociability and social novelty preference tests were conducted in a modified OF box to create a U-shaped two-choice test^30^. These tests evaluate two aspects of social behavior, including sociability, which compares the preference of mice to interact with social conspecifics or non-social objects, and social novelty, which assesses the tendency of mice to explore or interact with novel mice compared to familiar conspecifics. This behavioral assessment consists of three consecutive testing phases, including habituation, sociability, and social novelty sessions (Experimental design, **Figure 7L**). During the first session, experimental mice were habituated to the testing environment and were allowed to freely explore the empty modified U-shaped OF apparatus containing two identical empty wire grid cages for 10 min (habituation session). After the testing interval, experimental mice were removed from the testing environment and returned to a temporary cage. An unfamiliar stimulus mouse was placed in one out of two empty grid cages, then experimental mice were returned to the testing apparatus and allowed to freely explore the environment for 10 min (sociability session). Object or social target locations were randomized and counterbalanced between experimental mice and across treatment groups. Stimulus/testing mice were age-, sex-, and strain-matched. Experimental mice were able to investigate and interact with unfamiliar social targets in a wire grid container that permitted visual, olfactory, tactile, and auditory cues, while preventing direct physical contact. This was compared to the exploration of an identical empty wire grid container on the other side of the U-shaped OF apparatus. Following this session, experimental mice were returned to a temporary new cage, and a second stimulus mouse was placed in the remaining empty wire grid cage, becoming the “novel” unencountered social target. The prior stimulus mouse remained in the same wire grid cage during the third testing session and became the “familiar” social target. Experimental mice are returned to the testing environment for 10 min (social novelty session) and then returned to temporary cages until sociability and social preference tests have been completed for all group-housed mice.

Time spent exploring or investigating social or non-social targets was recorded and manually scored. Social investigation was defined as active or direct engagement with the subject mouse’s head oriented towards the wire grid cage within a 2 cm investigation zone, including sniffing, but excluding behaviors such as climbing or sitting on top of the wire grid cages. Locomotor activity patterns were tracked and recorded from video footage using ANYmaze behavior software. Total Investigation time (seconds) for either social or non-social targets was reported as a percentage of total investigation time (stimulus mouse + non-social object, or familiar social target), and preference indexes were calculated using the following formula: [investigation time with test mouse (s) / total investigation time]. Experimental mice were excluded from analyses if stimulus mice escaped from the wire grid cage or did not enter one of two zones during the testing session (sociability session, H5N1, n=2; social novelty preference session, Mock, n=2, and H5N1 = 2).

### Statistical Analyses

Statistical analyses were performed using GraphPad Prism v10.3.1 for Mac, GraphPad Software, Boston, Massachusetts, USA, www.graphpad.com. Kaplan-Meier curves were compared using the log-rank Mantel-Cox test. The area under the curve (AUC) of body mass change and rectal temperature curves was analyzed by a two-tailed unpaired t-test. Cumulative clinical scores were analyzed using the Kruskal-Wallis test. Viral titers, antibody measures, pup measurements, developmental, and behavioral tests were analyzed using two-tailed unpaired t-test or 1-way or 2-way ANOVAs followed by post hoc Tukey multiple comparisons test. Mean or median differences were considered statistically significant at P ≤ 0.05 and are represented by a single asterisk.

## Acknowledgments

We are grateful for the animal care staff from the Miller Research Building Animal Facilities at Johns Hopkins University. Light microscopy images were generated using the instruments and support of the Light Microscopy Core of the Department of Molecular Microbiology and Immunology at the Johns Hopkins Bloomberg School of Public Health. We also thank the histology laboratory personnel at the Johns Hopkins Oncology Tissue Services, and the Klein, Davis, and Thompson lab members for discussions. We also acknowledge Kimberly Davis for input on placental immunofluorescence images and Ryan Lewandowski for his technical assistance with digital histology scans at Michigan State University. Funding was provided by the National Institutes of Health, NIAID Johns Hopkins Center of Excellence for Influenza Research and Response (75N93021C00045; AP and SK), NIH/NIAID F32HD117584 (BAS), and the NIH/NIAID 5T32AI007417-29 (MAP).

## Author contributions

Conceptualization: BAS, MAP, AP, SLK

Methodology: BAS, LC, ASB, RJW, JRH, AP, SLK

Investigation: BAS, MAP, SNT, TZ, SC, JAL, JS, OH

Visualization: BAS, OH, JRH

Funding acquisition: SLK, AP

Project administration: SLK

Supervision: JRH, AP, SLK

Writing – original draft: BAS, JRH, AP, SLK

Writing – review & editing: All authors

