## Supplemental Files for "Trimester-Dependent Vertical Transmission of H5N1 Influenza Virus Through Placental and Mammary Routes Impairs Offspring Development"

**Supplemental Figures:**

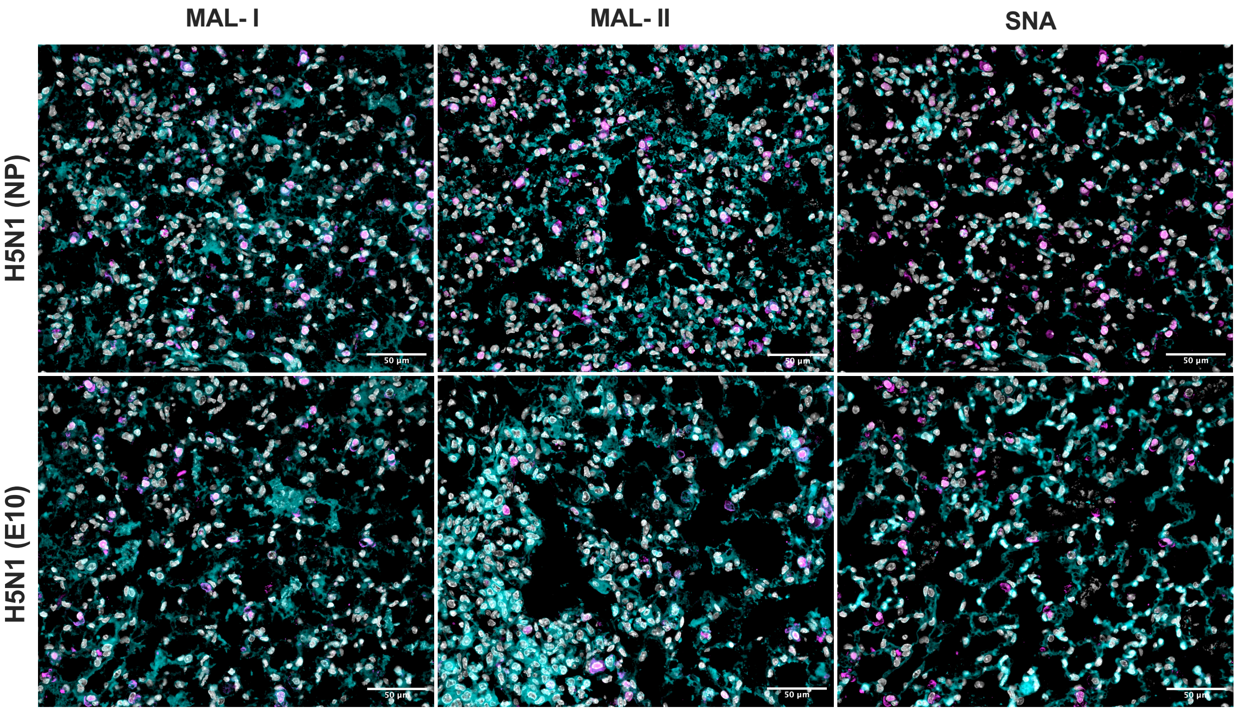


**Supplementary Fig. 1: Lectin and viral protein staining in the lungs of non-pregnant and pregnant female mice infected with bovine H5N1 influenza A virus.** CD-1 non-pregnant (7-8 weeks; n=5/group; NP) or pregnant (Embryonic day (E) 10; n=6-7/group) female mice were intranasally inoculated with 100 TCID50 of A/bovine/OH. At 6 days post-injection (dpi), mice from each group were euthanized, and their lung tissue was collected and fixed for immunofluorescent staining. Cell nuclei (DAPI) are labeled in grey, MAL-I (α 2,3 SA), MAL-II (α 2,3 SA), or SNA (α 2,6 SA) in cyan, and H5N1 virus protein in magenta. Images were captured at 20X (scale bar: 50 µm).

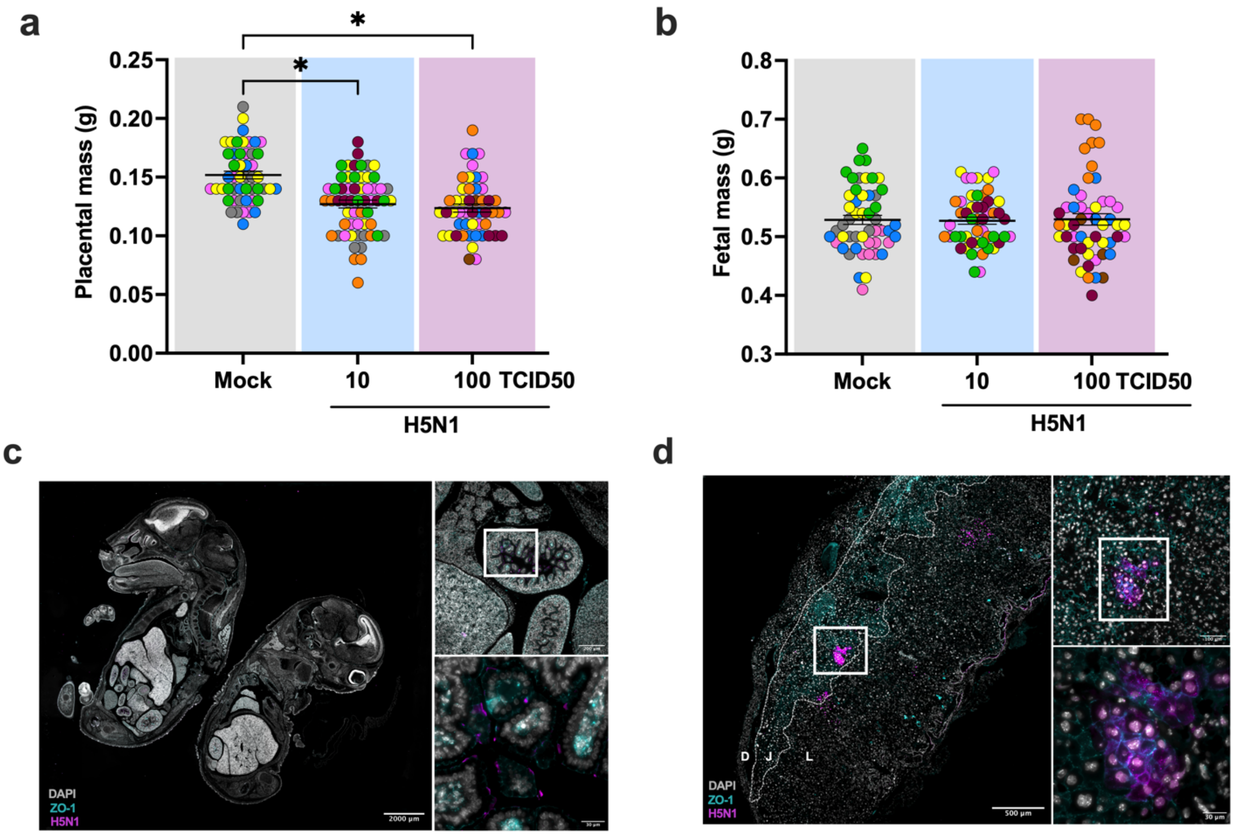


**Supplementary Fig. 2: Tissue tropism of bovine H5N1 influenza A virus infection during mid-gestation.** CD-1 pregnant females (E10) were intranasally inoculated with 10 or 100 TCID50 of A/bovine/OH. At 6 dpi, a subset of mice (n=5-7) from the dams inoculated with 10 TCID50 were euthanized and tissues were collected. The weight of (**a**) individual placentas and their (**b**) corresponding fetuses were recorded at the time of necropsy. Individual points are color-coded by litter, while shaded colors indicate group: grey for mock, blue for 10 TCID50, and purple for 100 TCID50 infected pregnant dams. Horizontal bars (B-C) indicate mean ± standard error of the mean (SEM) per group. Asterisks represent significant differences ($p < 0.05$) by a two-way ANOVA with Tukey post-hoc test. Immunofluorescent staining of the (**c**) fetus from E10 pregnant dams inoculated with H5N1. The left image was captured as a 4X tiled image (scale bar: 2,000 µm); higher-resolution images were captured at 10X (scale bar: 200 µm) and/or 60X (scale bar: 20 µm). White boxes indicate the regions displayed in the magnified images. Cell nuclei (DAPI) are shown in grey, tight junctions (ZO-1) in cyan to distinguish structural features, and H5N1 whole virus protein in magenta. Immunofluorescent staining of the (**d**) placenta from E10 pregnant dams inoculated with H5N1. The left image was captured at 4X (scale bar: 500 µm); higher-resolution images were captured at 20X (scale bar: 100 µm) and/or 60X (scale bar: 30 µm). White boxes indicate the regions displayed in the magnified images. Cell nuclei (DAPI) are shown in grey, tight junctions (ZO-1) in cyan to distinguish structural features, and H5N1 whole virus protein in magenta. The three zones of the mouse placenta (the decidua (D), junctional zone (J), and labyrinth (L)) are distinguished by the dashed white lines and labeled accordingly.

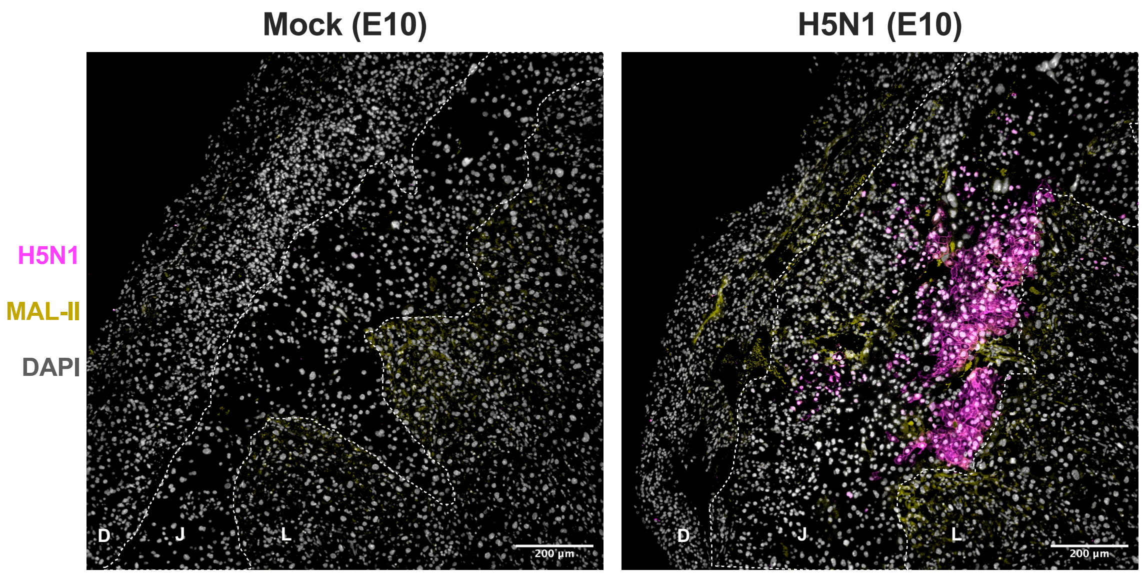


**Supplementary Fig. 3: Lectin and viral protein staining of bovine H5N1 influenza A virus within the junctional and labyrinth zones of the mouse placenta.** CD-1 dams (E10) were intranasally inoculated with media (Mock) or 10 TCID50 of A/bovine/OH (H5N1). At 6 dpi, dams (n=3-5/group) were euthanized and placentas (n=2-3/dam) were collected for immunofluorescent staining. Images were captured at 10X (scale bar: 200 µm). Immunofluorescence staining of MAL-II (α 2,3 SA) is shown in yellow, cell nuclei (DAPI) are shown in grey, and H5N1 whole virus antigen is shown in magenta. The three zones of the mouse placenta (the decidua (D), junctional zone (J), and labyrinth (L)) are distinguished by the dashed white lines and labeled accordingly.

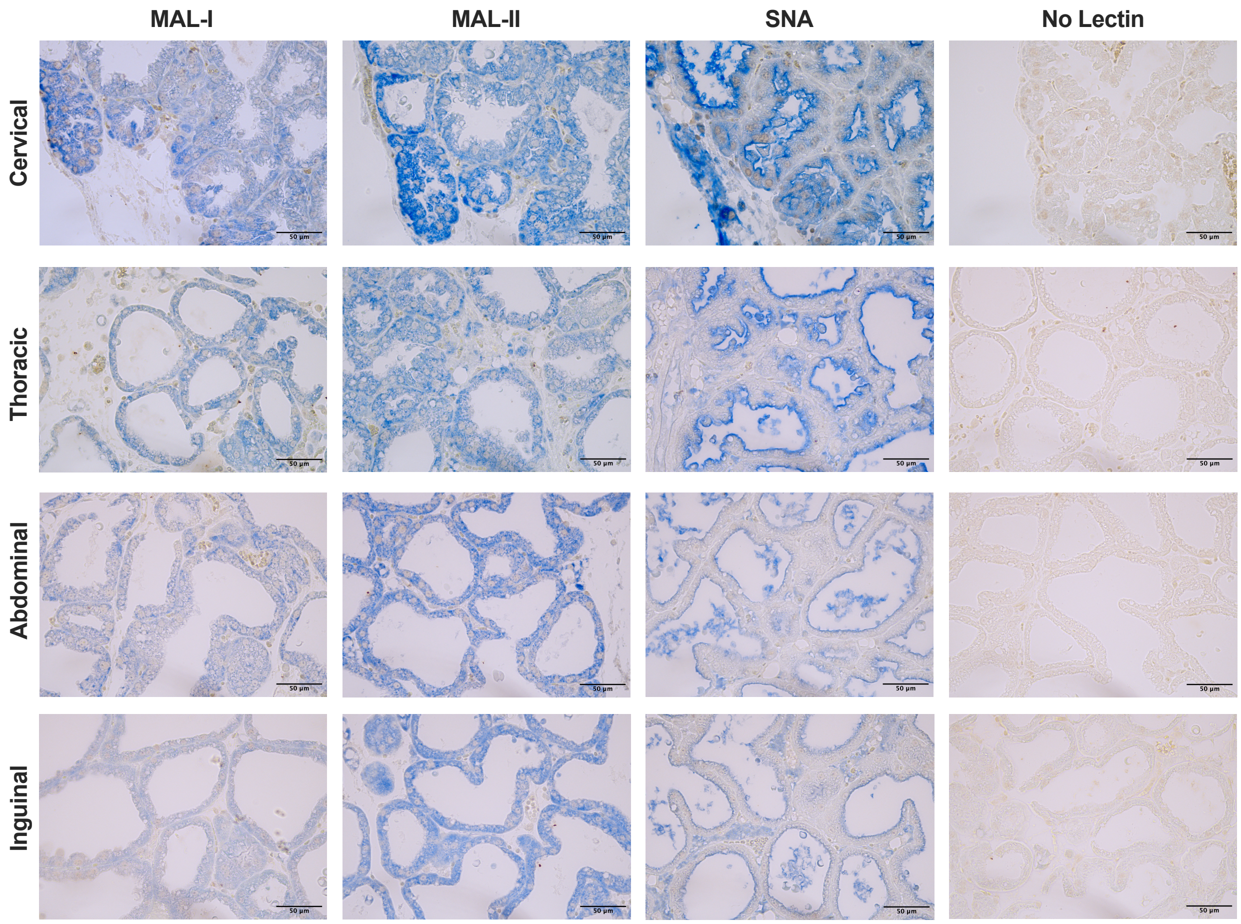


**Supplementary Fig. 4: Immunohistochemistry of mammary tissue from mock-infected E16 dams 4 days post-birth.** Cervical, thoracic, abdominal, and inguinal mammary tissue were collected 7-8 dpi from E16 pregnant females inoculated with media and fixed for histological analysis. Immunohistochemistry for viral N1 antigen (brown chromogen), individually duplexed with MAL-I (α 2,3 SA), MAL-II (α 2,3 SA), or SNA (α 2,6 SA) staining (blue chromogen). A control lacking the lectin primary antibody is also included. Images were captured at 40X (scale bar: 50 μm).

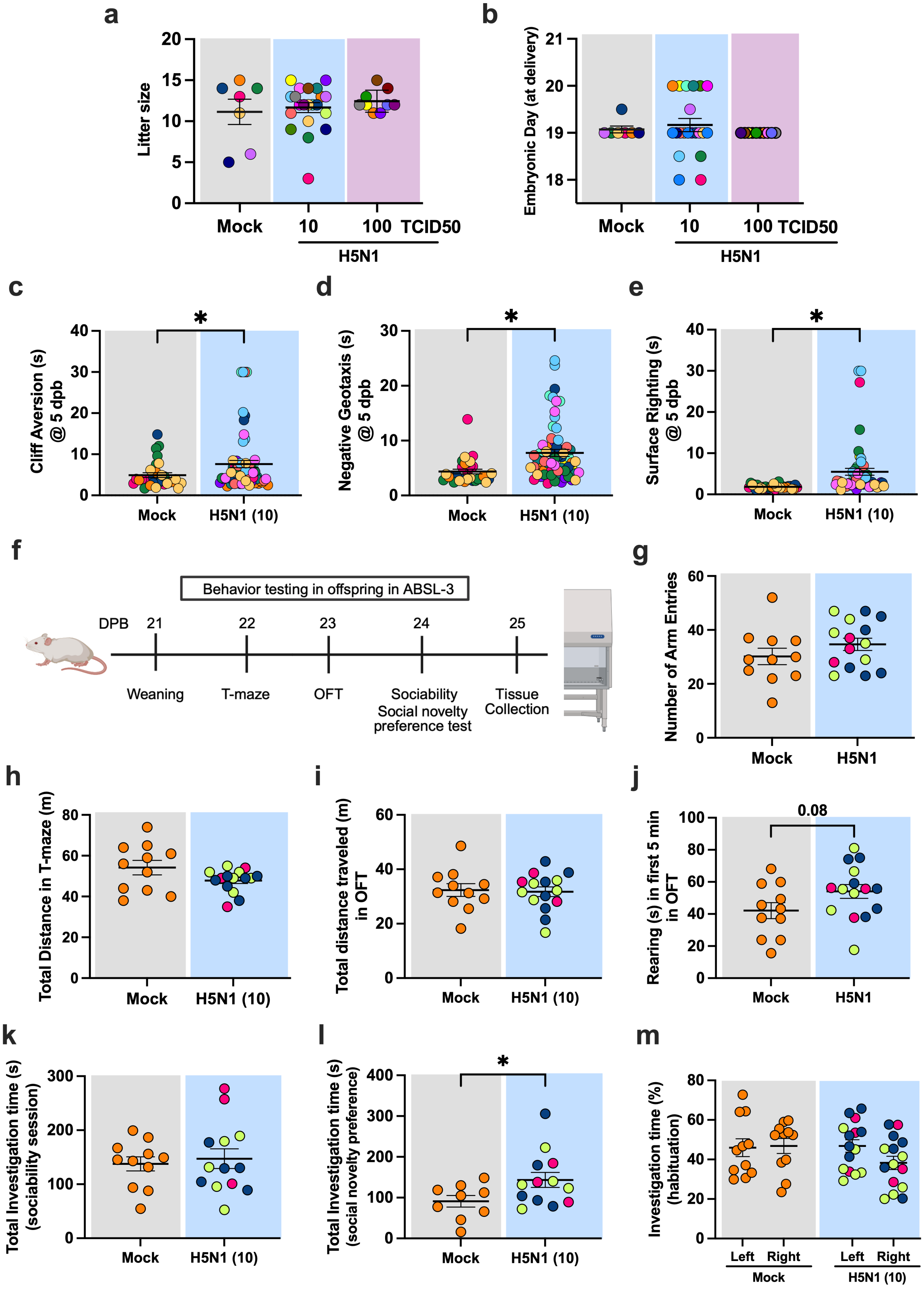

a
Litter size
Mock
10
100 TCID50
H5N1
b
Embryonic Day (at delivery)
Mock
10
100 TCID50
H5N1
c
Cliff Aversion (s) @ 5 dpb
*
Mock
H5N1 (10)
d
Negative Geotaxis (s) @ 5 dpb
*
Mock
H5N1 (10)
e
Surface Righting (s) @ 5 dpb
*
Mock
H5N1 (10)
f
Behavior testing in offspring in ABSL-3
DPB
21
22
23
24
25
Weaning
T-maze
OFT
Sociability
Social novelty preference test
Tissue Collection
g
Number of Arm Entries
Mock
H5N1
h
Total Distance in T-maze (m)
Mock
H5N1 (10)
i
Total distance traveled (m) in OFT
Mock
H5N1 (10)
j
Rearing (s) in first 5 min in OFT
0.08
Mock
H5N1
k
Total Investigation time (s) (sociability session)
Mock
H5N1 (10)
l
Total Investigation time (s) (social novelty preference)
*
Mock
H5N1 (10)
m
Investigation time (%) (habituation)
Left
Right
Mock
Left
Right
H5N1 (10)

**Supplementary Fig. 5: Perinatal outcomes following bovine H5N1 influenza A virus infection during third-trimester gestation.** CD-1 pregnant (E16) females were intranasally inoculated with 10 or 100 TCID50 of A/bovine/OH. (**a**) The litter size and (**b**) embryonic day of delivery were recorded. At 5 days post-birth (dpb) (**c-e**), a subset of pups from dams inoculated with media or 10 TCID50 of H5N1 from each litter was assessed for (**c**) cliff aversion, (**d**) negative geotaxis, and **(e)** surface righting to measure neurological development. (**f**) After weaning, behavioral phenotypes were analyzed using multiple cognitive assessments at 23 dpb, and social behavior at 24 dpb prior to tissue collection at 25 dpb. Spatial working memory was assessed in mice by measuring the (**g**) number of arm entries and (**h**) total distance travelled during the T-maze testing interval. (**l**) Overall locomotor activity was assessed by calculating the total distance traveled during the 10-minute test interval using the open field test (OFT). (**j**) The number of rearing bouts in the first 5 min of the OFT was quantified to examine anxiety-like behaviors. The sociability and social novelty preference test was used to assess preference for interacting with a novel or familiar mouse. The total investigation time was also calculated for the (**k**) sociability session and (**l**) social novelty preference session. (**m**) During the first testing session, termed habituation, the investigation time was measured for each side (left and right) of the modified U-shaped OFT apparatus. Individual points are color-coded by litter, while shaded colors indicate group: grey for media, blue for 10, and purple for 100 TCID50-inoculated dams. Horizontal bars indicate mean ± SEM per group. Asterisks represent significant differences ($p < 0.05$) by a (a-b, k) two-way ANOVA with Tukey post-hoc test or (c-m) two-tailed unpaired t-test.
